# A two-enzyme adaptive unit within bacterial folate metabolism

**DOI:** 10.1101/120006

**Authors:** Andrew F. Schober, Andrew D. Mathis, Christine Ingle, Junyoung O. Park, Li Chen, Joshua D. Rabinowitz, Ivan Junier, Olivier Rivoire, Kimberly A. Reynolds

## Abstract

Metabolic enzyme function and evolution is influenced by the larger context of a biochemical pathway – deleterious mutations or perturbations in one enzyme can often be compensated by mutations to others. To explore strategies for mapping adaptive dependencies between enzymes, we used a combination of comparative genomics and experiments to examine interactions with the model metabolic enzyme Dihydrofolate Reductase (DHFR). Biochemically, DHFR shares a metabolic intermediate with numerous folate metabolic enzymes. In contrast, comparative genomics analyses of synteny and gene co-occurrence indicate a sparse pattern of evolutionary couplings in which DHFR is coupled to the enzyme thymidylate synthase (TYMS), but is relatively independent from the rest of folate metabolism. To test this apparent modularity, we used quantitative growth rate measurements and forward evolution in *E. coli* to demonstrate that the two enzymes are coupled to one another, and can adapt independently from the remainder of the genome. Mechanistically, the coupling between DHFR and TYMS is driven by a constraint wherein TYMS activity must not greatly exceed that of DHFR – both to avoid depletion of reduced folates and prevent accumulation of the metabolic intermediate dihydrofolate. Extending our comparative genomics analyses genome-wide reveals over 200 gene pairs with statistical signatures similar to DHFR/TYMS, suggesting the possibility that cellular pathways might be decomposed into small adaptive units.

## Introduction

The collective action of enzymes in metabolism produces the basic materials for cells to grow and divide. Biochemical pathway maps provide a rich description of the reactions and intermediates needed for metabolism, yet it remains difficult to predict how metabolic systems will respond to perturbation. That is, if the activity or expression level of a particular enzyme were changed, which (if any) of the other enzymes in metabolism would require compensatory modification? Because the pattern of such adaptive interactions between proteins remains largely unknown, our ability to rationally engineer new systems [1–3], understand how the cell responds to perturbations [4, 5], and quantify the relationship between mutations and disease [6, 7] is limited. An ability to globally map dependencies between proteins and to identify groups of enzymes within a pathway that adapt as distinct subunits would help render cellular systems more tractable and predictable.

To begin to address this problem, we used a combination of comparative genomics and experiments to measure the pattern of adaptive interactions with the folate metabolic enzyme Dihydrofolate Reductase (DHFR). DHFR is an essential enzyme involved in the synthesis of purine nucleotides, thymidine and several amino acids [8]. As such, it is a common target of antibiotics, antimalarials, and chemotherapeutics [9, 10]. In recent years, it has become a prominent model system for understanding the evolution of drug resistance [11–15], evolution of protein conformational dynamics [16–18], and constraints on horizontal gene transfer [19, 20]. However, it remains unclear how changes to other enzymes in folate metabolism influence the relationship between DHFR function and cellular fitness. Understanding how mutations in other enzymes can compensate for loss of DHFR function is important to questions of antibiotic resistance and the evolution of folate metabolism. More generally, DHFR provides a well-studied model system to test computational approaches for predicting adaptive interactions.

Examining patterns of evolutionary coupling across species presents a practical strategy for inferring genes that share an adaptive constraint. We used analyses of both synteny (conservation of chromosomal proximity) and gene co-occurrence across 1445 bacterial genomes to create a statistical map of evolutionary couplings within folate metabolism. Though the folate biochemical pathway map is highly interconnected (many enzymes share a metabolic intermediate), our comparative genomics analyses suggest a sparse architecture of adaptive interactions in folate metabolism. Four small groups of genes appear as independent co-evolutionary units: the genes within a group co-evolve with each other, but each group appears to evolve relatively independently from the remainder of folate metabolism (and the genome). One of these gene groups contains the enzymes DHFR and Thymidylate Synthase (TYMS). Overall, these results suggest that small enzyme groups might adapt as relatively independent units inside the broader context of folate metabolism.

To test whether DHFR and TYMS are more coupled to each other than to the remainder of the biochemical pathway, we used quantitative measurements of growth rate, metabolomics and forward evolution in *E. coli* to map couplings to both DHFR and TYMS. We demonstrate that these two enzymes are (1) coupled through a shared metabolite, (2) are relatively less coupled to the remainder of folate metabolism and (3) can adapt independently from the remainder of the genome. In particular, we observe that mutations in these two enzymes are sufficient for *E. coli* to acquire resistance to the inhibition of DHFR by trimethoprim. Extending our statistical analyses of co-evolution genome wide reveals additional gene pairs that co-evolve with one another yet are relatively independent from the rest of the genome. This provides a set of hypotheses for future experimental testing, and suggests that the finding of small adaptive units embedded within larger cellular systems may be a general feature of cellular networks.

## Results

### A comparative genomics-based map of evolutionary coupling in folate metabolism

Starting from DHFR (encoded by the *E. coli* gene *folA*), we selected 16 folate metabolic genes for study using information from the STRING database (v10.5, [21]) and KEGG pathway maps [22]. STRINGdb is an online database of known and predicted protein interactions that uses a combination of information – including synteny, co-occurrence, online databases, co-expression, high-throughput experiments (e.g. yeast two hybrid data), and automated text mining – to produce an amalgamated confidence score [21]. This amalgamated score is benchmarked to predict the likelihood that two enzymes are in the same KEGG pathway. We selected all highest confidence interactions to DHFR (encoded by the *E. coli* gene *folA*). This results in fourteen genes, all of which encode an enzyme sharing a product or substrate with DHFR. We added two more genes (*metF, folD*) to this set to complete the folate cycle on the basis of KEGG pathway information.

This selection of genes encompasses core folate metabolism – a set of reactions that interconvert various folate species and produce methionine, glycine, thymidine and purine nucleotides in the process (Fig 1A and Table S1) [8]. The input of the pathway is 7,8-dihydrofolate (DHF), produced by the bifunctional enzyme dihydrofolate synthase/folylpolyglutamate synthetase (FPGS) through the addition of L-glutamate to dihydropteroate. Once DHF is formed, it is reduced to 5,6,7,8-tetrahydrofolate (THF) by the enzyme dihydrofolate reductase (DHFR) using NADPH as a co-factor. THF can then be modified by a diversity of one-carbon groups at both the N-5 and N-10 positions, and subsequently serves as a one-carbon donor in several critical reactions, including the synthesis of a few amino acids and purine nucleotides (Fig 1A, bottom square portion of pathway). The only step that oxidizes reduced folate (5,10-Methylene THF) back to DHF is catalyzed by the enzyme thymidylate synthase (TYMS), which modifies uridine monophosphate (dUMP) to thymidine monophosphate (dTMP) in the process.

**Figure 1.**
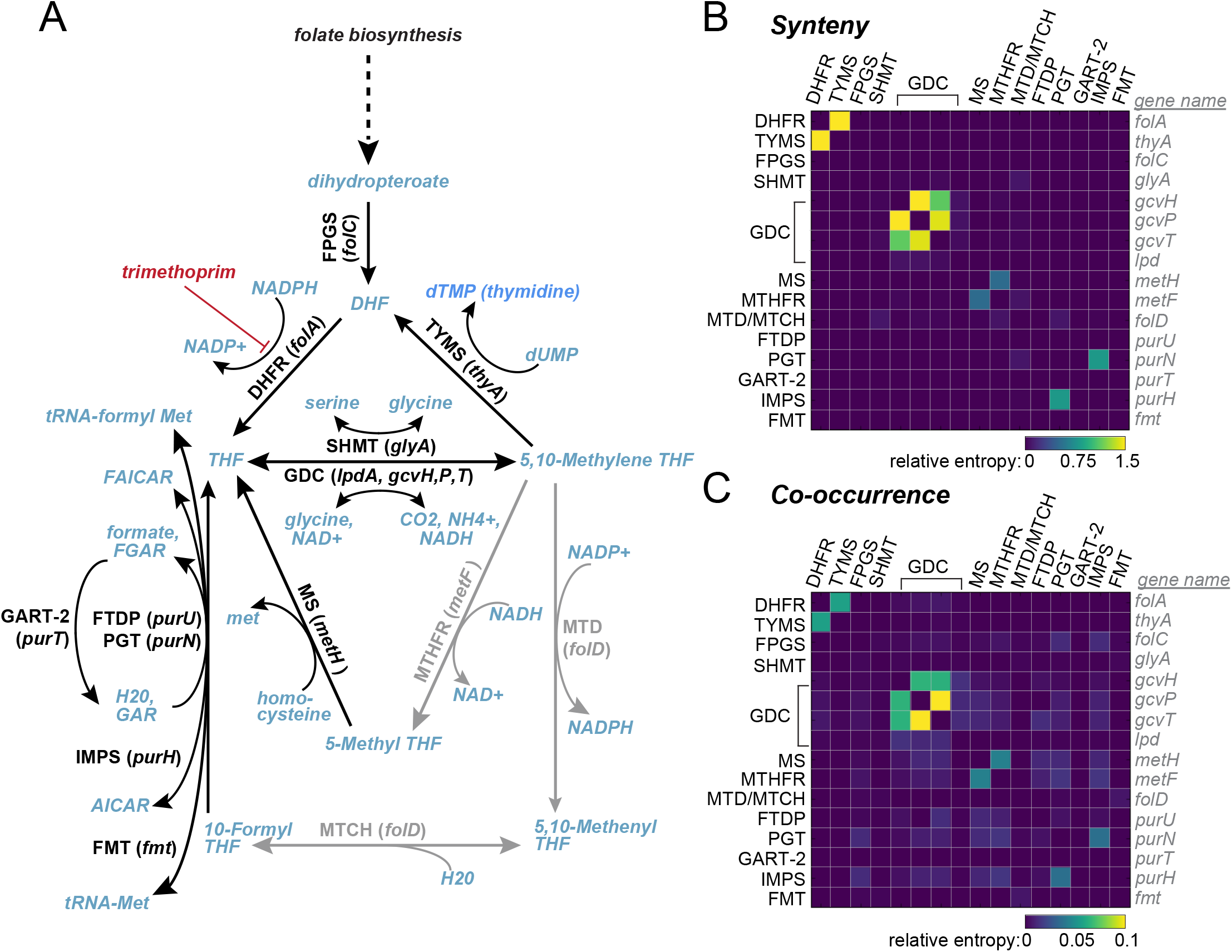
Biochemical and statistical representations of folate metabolism. **A,** Biochemical pathway map of folate metabolism. Blue font indicates metabolites, abbreviated enzyme (*gene*) names are in black or grey text. Black text and lines correspond to enzymes annotated as highest confidence interactions for DHFR (*folA*) in STRINGdb v10.5. See Table S1 for a more complete description of each enzyme. **B-C,** Heatmaps of evolutionary coupling between gene pairs in folate metabolism as evaluated by gene synteny and co-occurrence. Pixel color shows the extent of coupling between genes, indicated as a relative entropy, 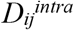. Enzyme names are indicated on the left and top of the matrix in black text, the corresponding gene names are given at right in grey italics. In *E. coli*, a single gene (*folD*) encodes a bifunctional enzyme that catalyzes both the methylene tetrahydrofolate dehydrogenase (MTD) and methenyltetrahydrofolate cyclohydrolase (MTCH) reactions in the biochemical pathway as shown in A. The majority of gene pairs show little co-evolution by either synteny or co-occurrence (dark purple pixels). See also Fig S1.

Consistent with the stated goals of the STRINGdb (to recover biochemical pathways), the STRING confidence scores indicate coupling between nearly all pairs of enzymes in folate metabolism. Both STRINGdb and the associated pathway map suggest that folate metabolism is highly interconnected, with many enzymes interacting via a shared product or substrate (Fig 1A, Fig S1A). To examine the extent to which these biochemical interactions lead to evolutionary couplings, we focused our analysis on measures that report on evolutionary selection across organisms. We computed two different statistical measures of evolutionary association across 1445 bacterial genomes: 1) synteny, the conservation of chromosomal proximity between genes and 2) co-occurrence, the coordinated loss and gain of genes across species [23–25]. Both approaches have been established as reliable indicators of protein functional relationships [24, 26–31], and technical variations on these methods are components of the amalgamated STRING score. These approaches are expected to report on any interaction important to evolutionary fitness, and can thus indicate gene pairs that interact through various mechanisms, including physical binding or biochemical constraints.

In contrast to the amalgamated STRING score, analyzing these two signals individually results in a sparse pattern of evolutionary couplings wherein most folate metabolic genes are relatively independent from each other (Fig 1B-C, Fig S1B-C). Consistent with expectation, we do observe evolutionary couplings between physically interacting gene products: the glycine cleavage system proteins H, P and T (*gcvH, gcvP*, and *gcvT* in *E. coli*) form a macromolecular complex [32]. We also see evolutionary couplings for three enzyme pairs: 1) DHFR/TYMS 2) methionine synthase (MS) with methionine tetrahydrofolate reductase (MTHFR) and 3) the purine biosynthesis proteins PGT and IMPS. While these enzyme pairs are not known to physically bind, each pair of enzymes share a metabolic intermediate, suggesting that a biochemical mechanism underlies their evolutionary coupling. Yet, most enzymes that catalyze neighboring reactions do not show statistical correlation (Fig 1, Fig S1). Instead, synteny and co-occurrence provide a different representation of folate metabolism in which most genes are relatively independent and a few interact to form evolutionarily constrained units.

So what factors might contribute to the observed sparsity of evolutionary couplings? One potential contributor is limited sensitivity of our methods. Indeed, false negatives – gene pairs that are evolutionarily coupled but without a strong synteny or co-occurrence signal – are expected due to limited statistical power. So perhaps the apparent modularity of the DHFR/TYMS pair within folate metabolism just reflects an incomplete pattern of predicted interactions. We find only six evolutionary couplings out of 120 possible pairwise interactions, even though 83 enzyme pairs interact biochemically (by sharing a product or substrate). Thus, a high false negative rate would be necessary to completely explain the difference between the pattern of biochemical interactions and the pattern of evolutionary couplings.

More generally, the pattern of adaptive interactions between enzymes need not strictly resemble the pattern of biochemical interactions, given the non-linear relationship between enzyme activities, metabolite concentrations, and fitness. The observed sparsity might then reflect a modular organization of adaptive constraints between enzymes, shaped by evolution. The evolutionary couplings are a statistical inference across thousands of genomes, and are not expected to capture idiosyncratic couplings specific to a particular organism or environmental condition. Instead, we hypothesize that the evolutionary coupling map presents predictions of the major adaptive couplings (and of the degree of independence). If true, this would suggest a route to begin decomposing metabolic pathways into meaningful, smaller multi-gene units, in a way that does not depend strongly on choice of model organism or environment.

As a first step towards testing this broader hypothesis, we set out to experimentally test if the statistical pattern of evolutionary coupling observed for DHFR and TYMS corresponds to the pattern of adaptive coupling in an individual organism, namely *E. coli*. In particular, are DHFR and TYMS indeed more coupled to each other than to other enzymes in the pathway, as the statistical results suggest? And can perturbations in TYMS suffice to compensate for perturbations in DHFR (without the need to modify other enzymes)? These proteins are not known to physically interact, providing an opportunity to better understand how biochemical constraints might drive co-evolution. Though the genes encoding DHFR and TYMS are proximal along the chromosome of many bacterial species, they are approximately 2.9 megabases apart in *E. coli*. Thus, our experiments will determine whether adaptive coupling between genes found in synteny persists even when they are distant on the chromosome.

### The growth rate defect of DHFR knockdown is partially rescued by changes in the expression of TYMS – but not other folate metabolic enzymes

As a first measurement of adaptive coupling between DHFR, TYMS and the rest of folate metabolism, we performed targeted measurements of growth rate dependency. More specifically, we decreased DHFR (or TYMS) expression, measured the effect on growth rate, and then examined if the effect on growth rate was modified in the background of a second gene knockdown. The null hypothesis is that the growth rate effect of reducing expression for one gene should be the same regardless of whether or not the second gene is knocked down – unless the two genes are coupled.

To make these measurements, we used CRISPR interference (CRISPRi) and a next-generation sequencing based measurement of relative growth rate. In CRISPRi, a single guide RNA (sgRNA) and a catalytically dead Cas9 enzyme are used to target and repress gene expression [33]. Pairs of sgRNAs can be cloned into a single vector, permitting knockdown of two genes at once (Fig S2A). Using this approach, we constructed pairwise knockdowns of DHFR and TYMS with every other gene in the folate metabolic pathway as described in Fig 1, with the exception of *lpdA* and *fmt*. We excluded *lpdA* and *fmt* from our experiments because they are involved in other critical cellular processes and are expected to have pleiotropic effects (*fmt* is necessary to produce the initiator methionine tRNA required for all protein translation; *lpdA* binds to the GDC but is also part of two other physical complexes in the cell). We also measured the effects of knocking down each gene individually by pairing each sgRNA with an sgRNA lacking the homology region needed for targeting a gene (referred to as none). In total, this results in a small library of 14 single and 25 double gene knockdowns (Table S2-S4). The knockdown activity of individual sgRNAs was verified by comparing the growth rate effect of knockdown to similar data for gene knockouts (data from the Keio collection, Fig S2C), and by qPCR (Fig S2D, Table S5).

To measure growth rate effects for the entire library, we transformed the library into MG1655 K12 *E. coli* and grew the cells in a turbidostat, a device for continuous culture that ensures: 1) the cells remain in exponential growth phase for the duration of the experiment 2) the media conditions are constant. We used M9 minimal media containing 0.4% glucose, 0.2% amicase, and 5 μg/ml thymidine for growth. These conditions were selected to be similar to previous forward evolution experiments ([15], with the addition of thymidine), and provide some measure of buffering for variation in the expression of folate metabolic enzymes. We took time points over the course of 12 hours, and then computed relative frequencies for each knockdown in the population by using next generation sequencing to “count” the number of each sgRNA pair in the population (Fig S2B). By fitting a line to the plot of relative frequency over time, we obtain a relative growth rate (slope) indicating the per hour decrease or increase in the abundance of a particular knockdown relative to wild type (carrying the targetless “*none*” sgRNA).

As expected, we observed that knocking down DHFR expression was highly deleterious. This was true whether alone or in combination with any other gene – with the exception of TYMS. Knocking down DHFR and TYMS together uniquely led to a partial rescue of growth rate (Fig 2A). Knocking down TYMS expression alone was also deleterious (though the effect is partly buffered by the presence of thymidine), and the deleterious effect was not improved by knocking down any other gene (Fig 2B). This indicates asymmetry in the coupling between DHFR and TYMS in this condition: knocking down TYMS buffers DHFR, but not vice versa. These results are consistent with prior attempts to generate *ΔfolA* strains of *E. coli*; when attempting to delete the gene encoding DHFR, a TYMS mutation is spontaneously acquired [34]. Overall the data show that among folate metabolic enzymes, decreasing TYMS expression uniquely compensates for reductions in DHFR expression.

**Figure 2.**
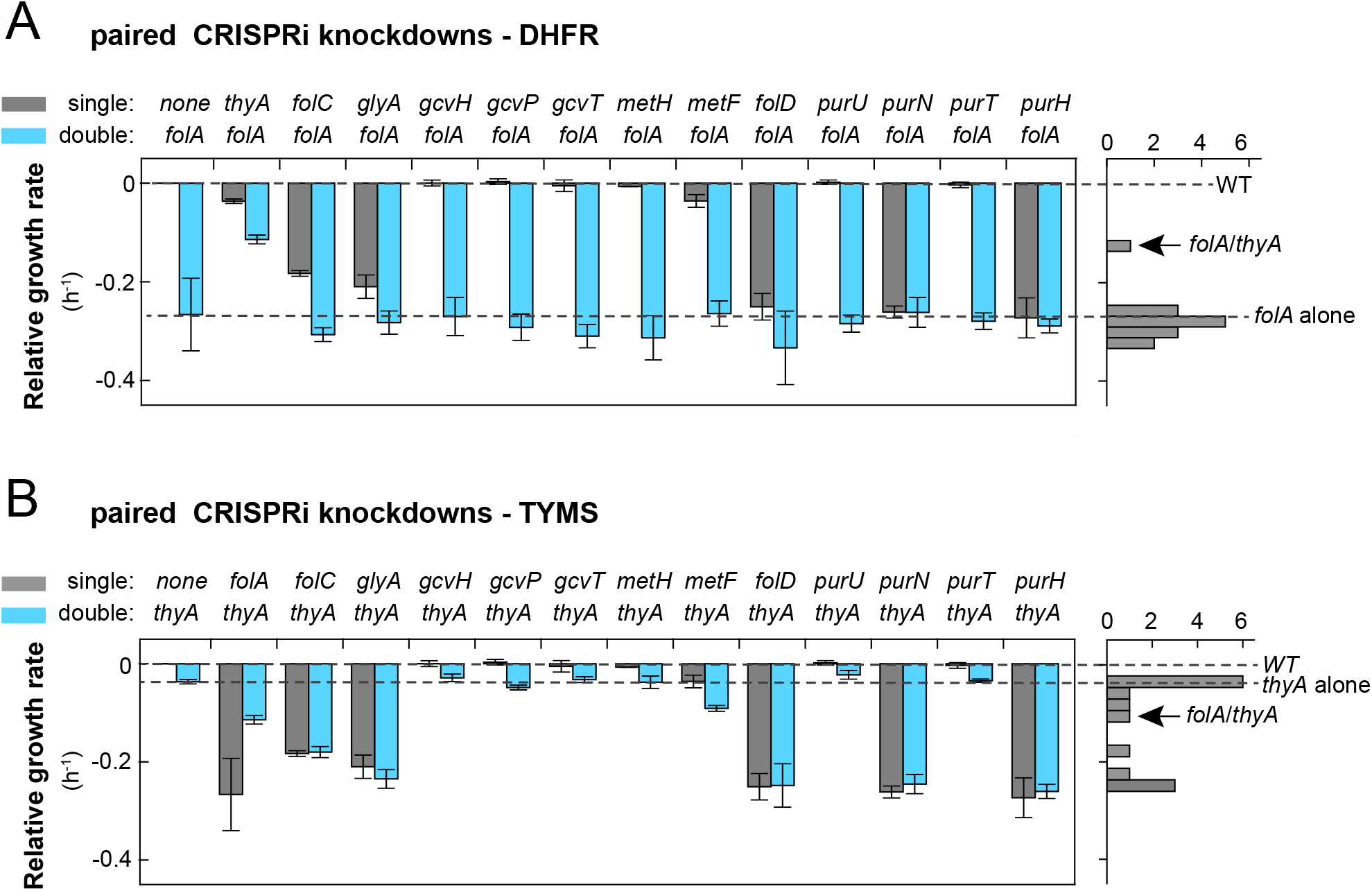
CRISPRi-based measurements of growth rate dependency for DHFR and TYMS. **A-B,** Relative growth rates for CRISPRi knockdowns of folate genes paired with either DHFR (**A**) or TYMS (**B**). Growth rates were measured in M9 minimal media supplemented with 0.2% amicase and 5μg/ml thymidine; and are calculated relative to the fitness of a strain carrying an sgRNA with no target homology region (none). Gray bars indicate the growth rate effect of a single mutant; blue bars indicate the effect of the double mutant. Error bars correspond to a 95% CI across at least three internal technical replicates (see also methods). The DHFR knockdown is significantly rescued by TYMS knockdown (*p*-value < 0.005 by Student’s t-test), but not by other gene knockdown (*p*-values > 0.20). As a reference point, the absolute doubling time of the *none* strain in turbidostat mono-culture is 0.83 ± 0.09 (95% CI) hours. From this, we estimate that a relative growth rate of −0.2 in these mixed population measurements corresponds to a doubling time of approximately 1.8 hours, and −0.4 is approximately zero growth rate (dead). To the right of each bar graph, we also plot a histogram of the growth rate effects for all knockdowns of DHFR (**A**) or TYMS (**B**).

### Forward evolution indicates adaptive independence of DHFR and TYMS from the rest of the genome

The above data are consistent with the idea that DHFR and TYMS are more adaptively coupled to each other than to other genes in folate metabolism, but leave open the possibility that DHFR and TYMS interact with other genes in the genome. Moreover, the comparative genomics analyses of synteny and co-occurrence suggest that the coupling between DHFR and TYMS leads them to co-evolve as a unit. To look for compensation by other enzymes genome-wide, and test the idea that DHFR and TYMS can adapt as an independent unit, we conducted a global suppressor screen. We inhibited DHFR with the common antibiotic trimethoprim and then examined the pattern of compensatory mutations with whole genome sequencing. If DHFR and TYMS indeed represent a quasi-independent adaptive unit, then suppressor mutations should be found within these two genes (*E. coli folA* and *thyA*) with minimal contribution from other sites.

To facilitate forward evolution in the presence of trimethoprim, we used a specialized device for continuous culture called a morbidostat [15, 35, 36]. The morbidostat dynamically adjusts trimethoprim concentration in response to bacterial growth rate and total optical density, thereby providing steady selective pressure as resistance levels increase (Fig S3A-C). The basic principle is that cells undergo regular dilutions with fresh media until they hit a target optical density (OD = 0.15); once this density is reached, they are diluted with media containing trimethoprim until growth rate is decreased. This approach makes it possible to obtain long trajectories of adaptive mutations in the genome with sustained phenotypic adaptation [15]. For example, in a single 13-day experiment, we observe resistance levels in our evolving bacterial populations that approach the trimethoprim solubility limit in minimal (M9) media.

As for the CRISPRi-based growth rate measurements, we conducted the forward evolution experiments in M9 minimal media supplemented with 0.2% amicase. We evolved three populations in parallel: in each case the media was supplemented with a different concentration of thymidine (5, 10, and 50 μg/ml) chosen to range from a condition in which TYMS loss-of-function is deleterious to one which fully rescues of TYMS activity. By alleviating selective pressure on the entire pathway, we sought to expose a larger range of adaptive mutations without biasing the pathway towards a particular result; this is similar to the common practice of conducting second site suppressor screens for essential genes under relatively permissive conditions [37]. In this context, an absence of mutation outside of the two-gene pair becomes more significant.

Over 13 days of evolution, we observed that the three populations steadily increased in trimethoprim resistance (Fig S3D, Fig S4). To identify the mutations underlying this phenotype, we selected 10 single colonies from the endpoint of each of the three experimental conditions for phenotypic and genotypic characterization (30 strains in total). For each strain, we measured the trimethoprim IC50, growth rate dependence on thymidine, and conducted whole genome sequencing (Fig 3, Fig S5 and Table S6-S8). All 30 strains were confirmed to be highly trimethoprim resistant (Fig 3, Table S6), with the exception of strains 4, 5 and 10 from the 50 μg/ml thymidine condition: these three strains grew very slowly at all concentrations of trimethoprim tested (up to 1500 μg/ml). Because the dependence of growth on trimethoprim did not fit a sigmoidal function for these three strains, we do not report an IC50. Further, strains from all three thymidine-supplemented conditions became dependent on exogenous thymidine for growth, indicating a loss of function in the *thyA* gene that encodes TYMS (Fig 3 and Fig S5). We confirmed that this loss of function is not a simple consequence of neutral genetic variation in the presence of thymidine; cells grown in 50 μg/ml thymidine and 0.2% amicase in the absence of trimethoprim retained TYMS function over similar time scales (Fig S6).

**Figure 3.**
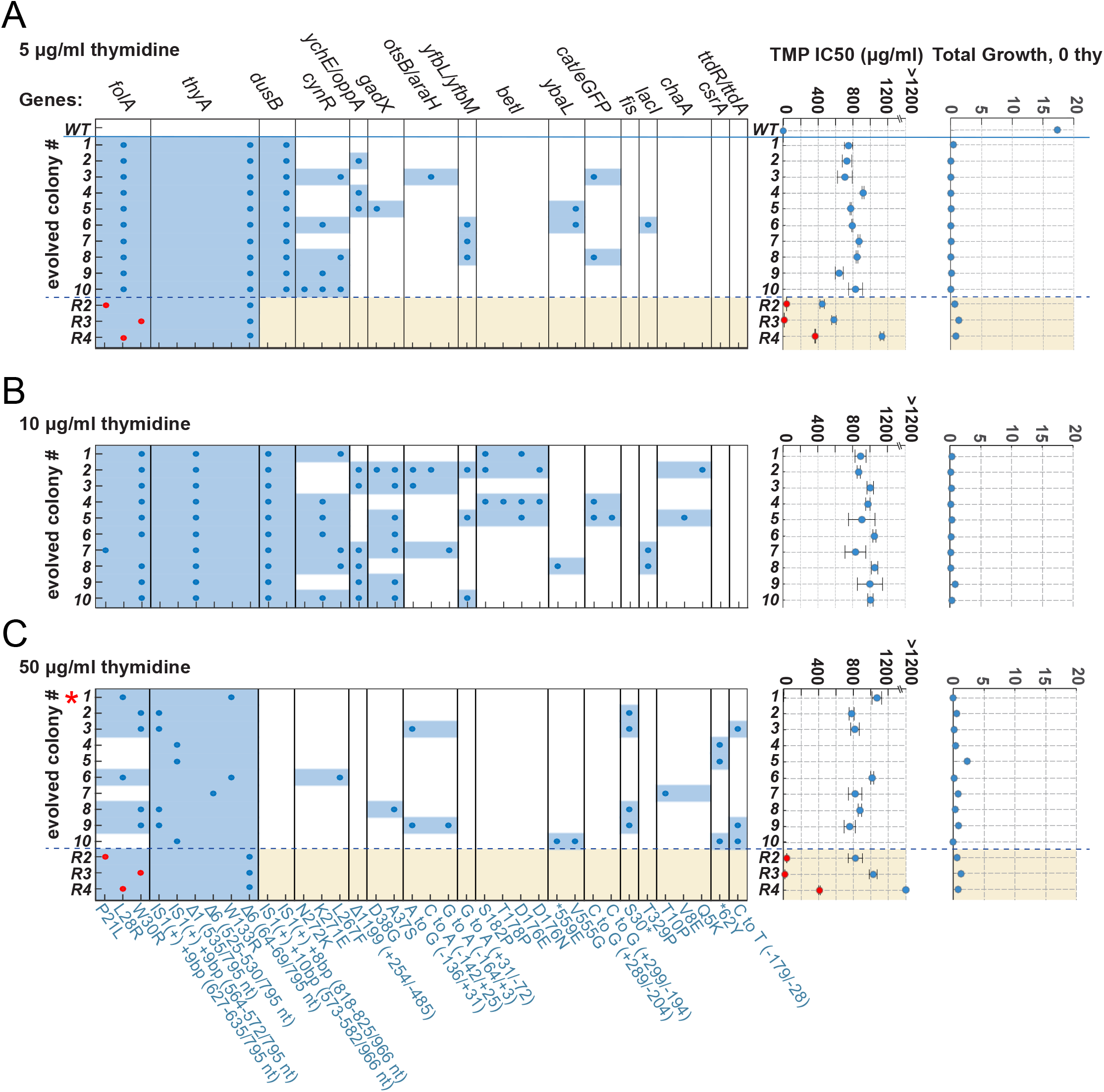
Genotype and phenotype of selected strains from three evolved populations. **A-C,** Ten single colonies (strains) were selected at the endpoint of each forward evolution condition for genotyping and phenotyping (30 in total). Panels A-C indicate the mutations observed in each strain sampled from that evolution condition. Genes that were mutated in two or fewer strains across all conditions are excluded, as are synonymous mutations (see Table S7 for sequencing statistics, and Table S8 for a complete list of mutants). Gene names are labeled along the top edge of the map, with the corresponding residue or nucleotide change(s) denoted along the bottom. If a strain acquires any mutation in a particular gene, the column section corresponding to that gene is shaded blue. All but four strains acquired mutations in both *folA* and *thyA*, encoding DHFR and TYMS, with the few exceptions lacking a *folA* mutation. A small red star indicates one strain with mutations in only DHFR and TYMS. To the right of each mutation map are trimethoprim (TMP) IC50 and thymidine dependence measurements for each strain. Error bars represent standard error over triplicate measurements. Evolved strains 4,5 and 10 (50 μg/ml thymidine) grew very slowly at all concentrations of trimethoprim measured, and we were unable to determine an IC50 by sigmoidal fit. Table S6 contains exact IC50 values and errors. Thymidine dependence is represented as area under the log(OD_600_) curve in 0 μg/ml thymidine over 10 hours. Evolved strains are no longer viable in the absence of extracellular thymidine, indicating a loss-of-function mutation in TYMS. See Fig S5 for growth rates across a range of thymidine concentrations. Three ‘reconstitution strains,’ featuring representative *folA* and *thyA* mutations recombined into a clean genetic background, have been included in panels A and C for comparison (denoted **R2-4**). Red dots in the rows of reconstituted genotypes R2-4 indicate the IC50 of a strain containing *only* the corresponding *folA* mutation. The *folA* single mutant strains were described in [13] and were produced through a different recombination protocol than the double mutants, and thus differ in the presence of additional chromosomal antibiotic markers but are otherwise identical.

Whole genome sequencing for all 30 strains revealed that isolates from all three conditions acquired coding-region mutations in both DHFR and TYMS, or even just in TYMS (Fig 3). The sequencing data for strains 4,5,7 and 10 (in 50 μg/ml thymidine) did not indicate any mutations or amplifications to DHFR or the promoter. For all other strains, the mutations identified in DHFR reproduce those observed in an earlier morbidostat study of trimethoprim resistance [15]. The mutations in TYMS – two insertion sequence elements, a frame shift mutation, loss of two codons, and a non-synonymous active site mutation – are consistent with loss of function. Thus, the observed pattern of mutation is consistent with our CRISPRi experiment: reduced TYMS activity can buffer inhibition of DHFR.

But are mutations in DHFR and TYMS sufficient to explain the resistance phenotype of the evolved strains? We did not observe mutations in any other folate metabolic genes during the course of the experiment, but we did observe that a few other genes mutated under multiple thymidine conditions (Table S9). These mutations might be neutral, contribute to trimethoprim resistance, or be adaptive for growth in continuous culture. Of particular interest were recurring mutations in *gadX*, a regulator of the acid response system, and the insertion sequences (IS1) in *dusB*. Trimethoprim is known to induce an acid response in *E. coli* that results in up-regulation of the *gadX* target genes *gadB/C* [38], and *dusB* is located in an operon upstream of the global E. coli transcriptional regulator *fis* [39]. Thus, we wanted to test if mutations in DHFR and TYMS alone were sufficient to explain the resistance phenotype. We introduced several of the observed DHFR and TYMS mutations into a clean wild-type *E. coli* MG1655 background (referred to as strains R1-R4) and measured the IC50. Strain R1 contained only a TYMS mutation (a deletion of six nucleotides, removing residues 25-26). As for strains 4,5 and 10 (in 50 μg/ml thymidine), strain R1 grew very slowly at all concentrations of trimethoprim measured (up to 1800 μg/ml), preventing determination of a reliable IC50. Strains containing mutations in both DHFR and TYMS (R2-R4) showed resistance phenotypes comparable to the evolved strains (Fig 3). For example, the DHFR L28R/ TYMS Δ6(64-69) double mutant in a clean genetic background (strain R4) was more resistant to trimethoprim than all of the evolved strains containing these two mutations (Fig 3A). For comparison, we also measured the IC50 for three previously constructed strains containing the DHFR single mutants (P21L, L28R, and W30R) [13]. The DHFR mutations alone were uniformly less trimethoprim resistant than the corresponding DHFR/TYMS double mutants. Taken together, we conclude that the paired DHFR and TYMS mutations are both necessary and sufficient to generate the observed resistance phenotype. Consistent with this, one of the evolved strains contains only mutations in DHFR and TYMS (strain 1 in 50 μg/ml thymidine, Fig 3C).

Interestingly, TYMS loss-of-function mutations have been observed in trimethoprim resistant clinical isolates for multiple genera of bacteria [40, 41]. This provides a degree of confidence that the laboratory evolution experiment reflects natural conditions of adaptation. Taken together the data support the idea that DHFR and TYMS represent a two-gene adaptive unit, with compensatory mutations contained within the unit.

### DHFR and TYMS are coupled by constraints on metabolite concentration

Given the above data, and no evidence for the physical association of DHFR and TYMS in bacteria, what is the mechanism underlying their adaptive coupling? The CRISPRi experiments and second site suppressor screen indicate that reductions in DHFR activity – either from reducing expression, or inhibiting catalysis – are buffered by decreases in TYMS activity. This suggests a simple hypothesis: their coupling arises from the need to balance the concentration of key metabolites in the folate metabolic pathway. In particular, both the depletion of reduced folates (THF species), and accumulation of DHF are expected to be detrimental to cell growth [42, 43]. So, we wanted to test the relationship between the activity of DHFR, the activity of TYMS, metabolite abundance and growth.

We started with a panel of ten DHFR mutations selected to span a range of *in vitro* catalytic activities [44], and measured the effect of these mutations on *E. coli* growth in the background of either WT TYMS or TYMS R166Q, a catalytically inactive variant. Again, we used a next-generation sequencing based assay to measure the relative fitness of all possible mutant combinations (20 total) [44]. In this system, DHFR and TYMS were expressed from a single plasmid that contains two DNA barcodes – one associated with DHFR and one with TYMS – that uniquely encode the identity of each mutant (Fig S7A). The full library of mutants was transformed into the thymidine auxotroph strain *E. coli* ER2566 *ΔfolA ΔthyA*, and grown as a mixed population in a turbidostat. Counting the frequency of the barcodes over time permits estimation of a relative growth rate for each mutant in the population (Fig S7B, Fig S8).

Relative growth rates were measured across two thymidine supplementation conditions: one in which TYMS R166Q produces a growth defect (5 μg/ml thymidine), and one that constitutes a full rescue of TYMS activity (50 μg/ml). In the background of WT TYMS (grey points, Fig 4A-B) growth rate was relatively insensitive to ~10-fold decrease in DHFR activity. As DHFR activity was diminished further, the relationship between catalytic power and growth rate became monotonic: decreases in DHFR activity result in slower growth. In the background of TYMS R166Q, decreases in DHFR catalytic activity were buffered. More precisely, we observed that the TYMS R166Q mutation is deleterious or neutral in the context of WT DHFR, but is beneficial in the context of low activity DHFR mutants (e.g. F31Y.G121V). In high thymidine conditions, this effect was very pronounced: the TYMS R166Q mutation was sufficient to rescue growth to WT-like levels for DHFR mutants with orders of magnitude less activity (Fig 4B).

**Figure 4.**
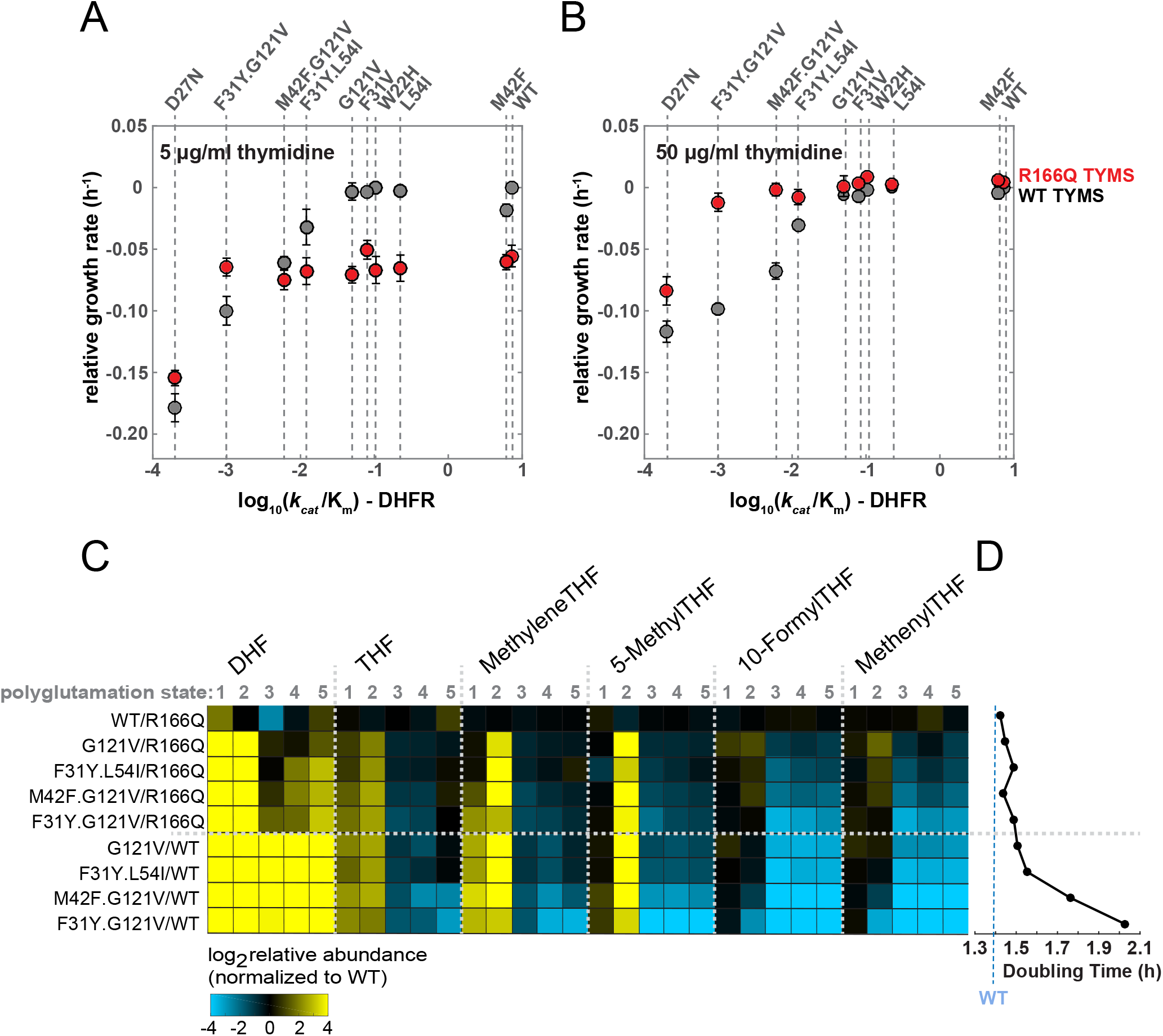
A loss-of-function mutation in TYMS buffers metabolic changes from decreased DHFR activity. **A, B**, Scatter plots of relative growth rate for DHFR mutants spanning a range of catalytic specificities (*k_cat_*/*K_m_*), and either a wild-type (WT, gray points) or catalytically dead (R166Q, red points) TYMS. Error bars correspond to standard error across triplicate measurements. In both conditions, we observe a buffering behavior in which the cost of reducing DHFR activity is mitigated by introducing a loss-of-function mutation in TYMS. **C,** Liquid chromatography-mass spectrometry profiling of intracellular folate species in M9 media supplemented with 0.1% amicase and 50 μg/ml thymidine. Rows reflect mutant DHFR/TYMS combinations, columns correspond to metabolites. Data represent the mean of three replicates, see also Fig S9 for associated errors. Each folate species can be modified by the addition of 1-5 glutamates. The color of each square denotes the log2 abundance of each species relative to wild-type. The data show that mutations reducing DHFR activity (G121V, F31Y.L54I, M42F.G121V, and F31Y.G121V) cause an accumulation of DHF and depletion of reduced folate species (THF) (bottom four rows). This effect is partly compensated by an inactivating mutation in TYMS (rows two-five). **D,** The corresponding doubling time for each mutant, as measured in batch culture (conditions identical to panel A).

From this set of mutants, we selected ten DHFR/TYMS pairs for liquid chromatography-mass spectrometry (LC-MS) profiling of folate pathway metabolites. The experiment was carried out for log-phase cultures in M9 glucose media supplemented with 0.1% amicase and 50 μg/ml thymidine, conditions in which the selected DHFR mutations displayed significant growth defects individually, but in which the corresponding DHFR/TYMS double mutants are restored to near wild-type growth. Current mass spectrometry methods allow discrimination between the full diversity of folate species, which differ in oxidation, one-carbon modification, and polyglutamylation states, permitting a broad metabolic study of the effects of mutations [45].

The data confirm that for DHFR loss-of-function mutants, intracellular DHF concentration increases (Fig 4C, bottom four rows). In addition, we observe depletion of reduced polyglutamated folates (Glu >= 3), while several mono- and di-glutamated THF species accumulate (particularly for THF, Methylene THF and 5-Methyl THF). Prior work found that DHF accumulation results in inhibition of the upstream enzyme FP-*γ*-GS [43]. FP-*γ*-GS catalyzes the polyglutamylation of reduced folates, an important modification that increases folate retention in the cell and promotes the use of reduced folates as substrates in a number of downstream reactions [46]. The observed pattern of changes in the reduced folate pool (depletion of polyglutamated THF forms) is consistent with inhibition of FP-*γ*-GS by DHF.

In the background of the corresponding TYMS loss-of-function mutant, the metabolite profiles become more WT-like: the depletion of reduced folates is less severe, and the accumulation of DHF more moderate (Fig 4C). Overall, the changes in metabolite concentration are consistent with the observed growth rate defects in the DHFR loss-of-function mutants: mutants with more severe DHF accumulation and THF depletion grow more slowly (Fig 4D, Fig S9). Thus, coordinated decreases in the activity of DHFR and TYMS serve to maintain balance in the DHF and THF metabolite pools, a condition associated with optimal growth. This provides a plausible biochemical mechanism for the strong statistical association of DHFR and TYMS by synteny and co-occurrence across thousands of bacteria. In this sense, the bifunctional fused form of DHFR/TYMS found in protists and plants could be regarded as an extreme case that guarantees stoichiometric expression [47, 48].

### Genome-wide analyses of co-evolution identifies additional adaptive pairs

The comparative genomics analyses of folate metabolism indicate a sparse pattern of adaptive interactions, in which DHFR and TYMS were more coupled to each other and less coupled to the remainder of folate metabolism. The experimental data support the interpretation that they form an adaptive unit embedded in folate metabolism: the two enzymes adapt together in response to trimethoprim, and they are linked by a shared constraint on metabolite concentration. Taken together, this suggests the hypothesis that analyses of synteny and gene co-occurrence might be used to identify small adaptive units elsewhere in the cell.

With this goal, we extended our analyses of gene synteny and co-occurrence to include all genes represented in *E. coli*. To ensure good statistics, we filtered the orthologs analyzed to those that co-occur in a sufficiently large number of genomes (2095 COGs, ~500,000 pairs in total) (see also Supplemental Experimental Procedures). We also compared our analysis to: (1) metabolic annotations from KEGG [22] and (2) the set of high-confidence binding interactions in *E. coli* reported by the STRINGdb [21]. Consistent with intuition and prior work, co-evolving gene pairs show enrichment for physical complexes, enzymes in the same metabolic pathway, and more specifically, enzymes with a shared metabolite (Fig 5A,B).

**Figure 5.**
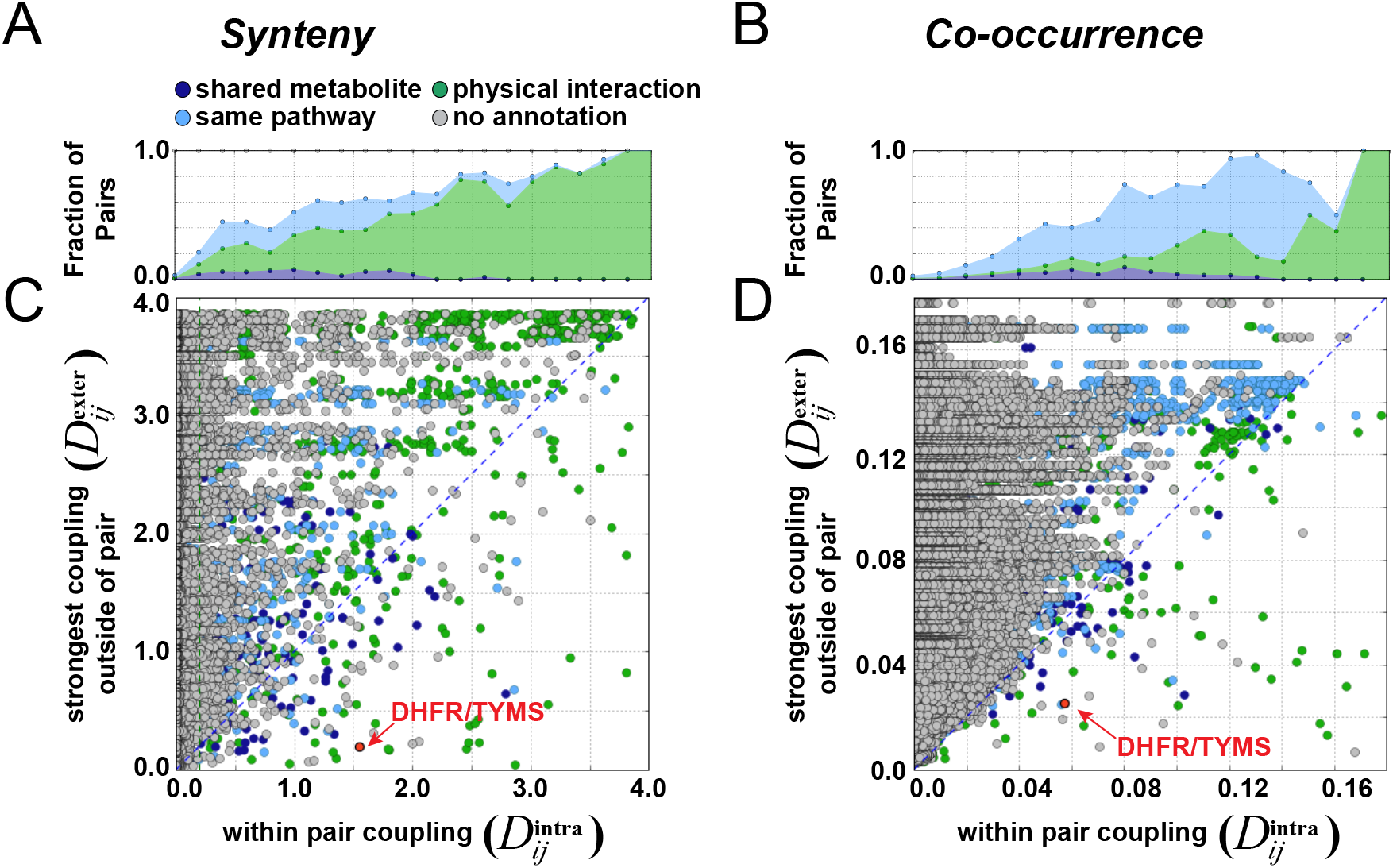
Genome-wide analysis of co-evolution in *E. coli*. **A,** Enrichment of physical and metabolic interactions as a function of synteny coupling. **B**, Enrichment of physical and metabolic interactions as a function of co-occurrence coupling. **C,** A scatterplot of synteny-based coupling for all analyzed gene pairs. Each point represents a pair of orthologous genes (COGs); coupling within the pair is shown on the x-axis, and the strongest coupling outside of the pair is shown on the y-axis. Color-coding reflects annotations from the STRING database (physical interactions) or KEGG database (metabolic pathways): green indicates binding, while pairs in dark blue or light blue are not annotated as physical interactions but are found in the same metabolic pathway. Dark blue gene pairs share a metabolic intermediate. The DHFR/TYMS pair is highlighted in red. See Table S10 for an annotated list of gene pairs below the diagonal. **D**, Scatterplot of coupling by co-occurrence for all analyzed gene pairs (same format as C).

To further identify pairs of genes that are strongly evolutionarily coupled to each other and relatively decoupled from the remainder of the genome, we constructed scatterplots of all gene pairs (Fig 5C,D). These plots indicate the strength of coupling within each pair (as a relative entropy, along the x-axis) versus the strongest coupling outside of the pair (along the y-axis). This analysis is limited to the identification of two gene units, and our analysis will need to be extended to identify larger communities within the complete network of co-evolutionary relationships [49]. It presents a simple graphical method for identifying gene pairs that are more tightly coupled to each other than any other gene in the dataset: these are the set of points below the diagonal (Table S10). Thus, these points represent predictions of two-gene coevolving units (258 by synteny, 194 by co-occurrence), and include the DHFR/TYMS pair. Many of the predicted adaptive units are two-protein physical complexes, while others share a metabolic intermediate like DHFR/TYMS.

In this plot, distance from the diagonal indicates the degree of modularity of a pair. By construction, points near the diagonal have an interaction with at least one other gene that is similar to the degree of coupling within the pair. The most modular pairs occupy the space in the lower right corner of the graph – by this criteria, DHFR and TYMS show an exceptionally strong signature of modularity. Considering the regime where 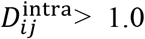 and 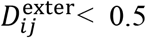 on the synteny plot, we observe a few other pairs with experimental evidence for adaptive coupling. For example the gene pair *accB/accC* which encodes two of the four subunits of acetyl-CoA carboxylase, the first enzymatic step in fatty acid biosynthesis. Overexpression of either *accB* or *accC* individually causes reductions in fatty acid biosynthesis, but overexpressing the two genes in stoichiometric amounts rescues this defect [50, 51]. Constraints on relative expression have also been noted for the *selA/selB* and *tatB/tatC* gene pairs [52, 53]. The *tatB/tatC* genes encode components of the TatABCE twin-arginine translocation complex, while *selA/selB* are not known to bind but are both involved in selenoprotein biosynthesis. The full set of gene pairs below the diagonal now serve as a starting place for more deeply understanding the hierarchical pattern of evolutionary couplings inside of cellular pathways, and testing the prevalence of adaptive modules more generally (Table S10).

### Discussion and Conclusions

Comparative genomics analyses of synteny and co-occurrence are well-established tools for the prediction of interactions between genes [24, 26–30]. However, they are not typically used to infer independence. Here we effectively ask whether, for evolutionary couplings between genes, the absence of evidence can be evidence for absence. Conceptually, this is analogous to recent work in proteins, in which co-evolution was used to map quasi-independent adaptive and functional residue groups within a single protein [54]. In our case study of folate metabolism, comparative genomics suggested a sparse pattern of evolutionary coupling between folate metabolic enzymes, with DHFR and TYMS showing significant coupling to each other but less coupling to the remainder of the folate metabolic pathway. Experimental analyses support the interpretation that these two enzymes form a relatively independent adaptive unit: they are more coupled to each other than any other enzyme in the pathway, they adapt together in response trimethoprim, and their activities are coupled by a shared constraint on metabolite concentrations. So, in this case, comparative genomics identified a meaningful adaptive unit much smaller than the scale of the entire biochemical pathway. In light of these findings, it becomes less obvious that shared membership in a KEGG metabolic pathway should be a default gold standard for interaction prediction approaches; instead the goals of the method and type of interaction being predicted (shared biochemical intermediate, physical complex or adaptive interaction) should be carefully considered.

DHFR and TYMS represent a first case study, and comparative genomics approaches are not expected to predict all possible adaptive interactions. Analyses of synteny and co-occurrence are an inference across thousands of species, encapsulating hundreds of millions of years of evolutionary divergence. In the context of specific environmental conditions (including conditions never encountered in past evolutionary history), or other model organisms, additional adaptive interactions likely exist. We expect that the adaptive units predicted by synteny and co-occurrence represent well-conserved, core, adaptive interactions that can sometimes be elaborated on under particular environmental conditions or in particular species. We propose that if a perturbation is made in one gene of the adaptive unit, the first, and most common adaptive mutations will also occur within the unit (either within the same gene, or in the other gene in the pair). Further, we expect that mutations within the pair should largely suffice to restore function, with mutations outside the pair having more subtle and/or idiosyncratic (e.g. environment or species specific) effects.

If it is generally possible to decompose metabolic pathways into smaller, relatively independent, pieces, this would suggest a route to identify units for biosynthetic engineering, and provide fundamental insights into how cells maintain homeostasis and adapt in the face of changing environments. For example, thymidine synthesis is the rate-limiting step for DNA synthesis in eukaryotic cells, and transcription of the TYMS and DHFR genes is greatly upregulated (via a common transcription factor) at the G1/S cell cycle transition [56]. In computational models of eukaryotic folate metabolism, computationally increasing the activities of DHFR and TYMS 100-fold results in increased thymidine synthesis but only modestly changes the concentration of other folates. Thus decoupling the DHFR/TYMS pair from the remainder of folate metabolism may be important for ensuring modularity in different metabolite pools. Substantial further work is needed to go beyond the DHFR/TYMS pair – to comprehensively test the relationship between the units identified by comparative genomics, patterns of adaptive dependencies, and adaptation to environmental changes. The data presented here provide the computational hypotheses, technical approaches, and motivation for now doing so.

## Materials and Methods

### Comparative genomics analyses

Synteny analysis was conducted using a simplified version of the methods described in [23, 24]. See the Supplemental Experimental Procedures for a detailed description of the synteny and co-occurrence calculations.

### CRISPRi-based growth rate measurements

We used next-generation sequencing based measurements of relative growth rate to quantify the effects of CRISPRi knockdowns on *E. coli* growth. The methods are described in detail in the Supplemental Information; here we provide a brief summary of the approach. Following from Qi et al., pairs of single guide RNAs (sgRNAs) were ligated into expression vectors using a single golden gate cloning reaction [33, 57]. We modified the sgRNA-containing plasmid to contain unique barcodes that allow for replicate measurements and error correction within a single experiment (see SI for more details). The resulting pairwise sgRNA library was transformed into MG1655 *E. coli* engineered to contain a chromosomally encoded, catalytically dead Cas9 (dCas9) under control of an inducible Tet promoter. The resulting transformants were grown in a continuous culture device (turbidostat) in M9 minimal media with 0.4% glucose, 35 μg/mL Kanamycin, 5 μg/mL thymidine, and 0.2% amicase for selection. After adaptation to the turbidostat, dCas9 was induced by the addition of anhydrous tetracycline (aTC) and culture time points were taken over 12 hours. From each time point, the sgRNA-containing region of each vector was deep sequenced to find the relative frequency of each sgRNA pair compared to a negative control sgRNA (lacking a homology targeting region). These data were used to calculate relative growth rates.

### Forward evolution of trimethoprim resistance in the morbidostat

The morbidostat/turbidostat apparatus was constructed as described by Toprak and colleagues [36]. The founder strain for the forward evolution experiment was *E. coli* MG1655 modified by phage transduction to encode green fluorescent protein (*egfp*) and chloramphenicol resistance (*cat*) at the P21 attachment site. The goal of this modification was to prevent and detect contamination with other strains. Throughout the forward evolution experiment, cells were grown at 30°C in M9 media supplemented with 0.4% glucose and 0.2% amicase (Sigma); 30 μg/ml of chloramphenicol (Cam) was added for positive selection.

To begin the experiment, the founder strain was cultured overnight at 37°C in Luria Broth (LB) + 30 μg/ml Cam. This culture was washed twice with M9, and back diluted into M9 + 30 μg/ml Cam supplemented with 5, 10, or 50 μg/ml thymidine (thy) for overnight adaptation in culture tubes at 30°C. The next day (henceforth referred to as day 0; day 1 is the end of the first day of adaptation), these overnight cultures were streaked onto LB agar plates: two colonies per condition were chosen for whole genome sequencing (WGS) in order to obtain an accurate sequence for the founder strain. The remainder of the overnight cultures was used to inoculate three morbidostat tubes at containing M9 media with varying thymidine supplementation (5, 10, and 50μg/ml thy). The starting optical density was approximately 0.005. Initial antibiotic concentrations were 0, 11.5 and 57.5 μg/ml trimethoprim for media stocks A, B, and C respectively. Each culture grew unperturbed until it surpassed an OD_600_ of 0.06, at which point it underwent periodic dilutions with fresh media. The dilution rate is given by the formula 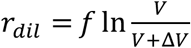, where *V* = = 15*ml* is the culture volume, and Δ*V* = 3*ml* is volume added. We chose a dilution frequency *f* = 3 h^-1^, to give *r_dil_* = 0.55. Above the OD_600_ = 0.15, these dilutions are used to introduce TMP into the culture (see also Fig S3). This allows controlled inhibition of DHFR activity in response to growth rate. Cycles of growth and dilution continued for a period of ~22 hours, at which point the run was paused to make glycerol stocks, replenish media, and update TMP stock concentrations. Culture vials for the next day of evolution were filled with fresh media and inoculated using 300μl from the previous culture. Complete trajectories of OD_600_ and drug concentration are shown in Fig S4. Endpoint cultures were streaked onto LB agar plates supplemented with 30 μg/ml of Cam and 50 μg/ml thymidine to obtain isolated colonies for whole genome sequencing.

### Whole genome sequencing

Two isolates were selected from each adapted day 0 culture, and ten clonal isolates (colonies) were randomly selected from the endpoint of each evolution condition, totaling 36 strains. Isolation of genomic DNA was performed using the QIAamp DNA Mini Kit (Qiagen). The Nextera XT DNA Library Prep Kit (Illumina) was used to fragment and label each genome for paired-end sequencing using a v2 300-cycle MiSeq kit (Illumina). Average read length and coverage can be found in Table S7. Genome assembly and mutation prediction was performed using *breseq* [58]. The reference sequence was a modification of the *E. coli* MG1655 complete genome (accession no. NC_000193), edited to include the GFP marker and chloramphenicol resistance cassette in our founder strain. The modified reference sequence and all complete genome sequences from the beginning and endpoint of forward evolution are available in the NCBI BioProject database (accession number: PRJNA378892).

### Measurements of thymidine dependence

All strains were grown overnight in LB + 5μg/ml thy, with the exception of the strains evolved in the 50μg/ml thy, which were supplemented with 50μg/ml thy to ensure viability. Cultures were then washed twice in M9 media without thymidine, and inoculated at an OD_600_=0.005 in 96-well plates containing M9 media supplemented with 10-fold serial dilutions of thymidine, ranging from 0.005 μg/ml to 50 μg/ml (in singlicate). OD_600_ was monitored in a Victor X3 plate reader at 30°C over a period of 20 hours. Growth was quantified using the positive integral of OD_600_ over time. This measure captures mutational or drug-induced changes in the duration of lag phase as well as perturbations in growth rate [15]. For each strain, we identified a start-time (*t*_0_) at the end of lag-phase for the fully-rescued 50μg/ml thy condition. We chose each *t*_0_ computationally as the last point before monotonic growth above the limit of detection. The log(OD_600_) versus time curves for all conditions are then vertically shifted (‘background-subtracted’), such that the function value at this start-time is zero. This curve is then numerically integrated from *t*_0_ to *t*_0_+15 hours using the trapezoid method.

### Measurements of trimethoprim resistance (IC50)

All strains were grown overnight in LB + 5μg/ml thy, with the exception of the strains evolved in the 50μg/ml thy, which were supplemented with 50μg/ml thy to ensure viability. Each strain was then washed into media conditions corresponding to the strain’s forward evolution condition, and adapted for 5.5 hours at 30°C. The recovery cultures were used to inoculate 96-well plates containing M9 media sampling serial dilutions of TMP (in triplicate), with a starting OD_600_ = 0.005. OD_600_ was monitored using a Tecan Infinite M200 Pro microplate reader and Freedom Evo robot at 30°C over a period of at least 12 hours. The trimethoprim resistance of each strain was quantified by its absolute IC50, the drug concentration (μg/ml) at which growth is half-maximal. The relationship between growth and trimethoprim inhibition is modeled using the four parameter logistic function:

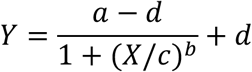

where *Y* is growth, *X* is TMP concentration, *a* is the asymptote for uninhibited growth, *d* is the limit for inhibited growth, *c* provides the concentration midway between *a* and *d*, and *b* captures sensitivity [59]. Growth was quantified using the positive integral of OD_600_ data over a 10h period of growth (see also the methods for measurement of thymidine dependence). For each strain, we identify a start-time (*t*_0_) at the end of lag-phase for the uninhibited 0μg/ml TMP condition. Growth versus TMP concentration was fit to the above model using MATLAB. IC50 was calculated as the concentration *X*\* for which growth *F*(*X*\*) = *a*/2.

### Growth without trimethoprim selection in 50μg/ml thymidine using the turbidostat

The founder strain for this experiment was identical to that used for evolution of trimethoprim resistance. Throughout the experiment, cells were grown at 30°C in M9 media supplemented with 0.4% glucose and 0.2% amicase (Sigma); 30 μg/ml of chloramphenicol (CAM) was added for positive selection. To begin the experiment, the founder strain was cultured overnight at 37°C in Luria Broth (LB) + 30 μg/ml Cam. This culture was washed twice with M9, and back diluted into M9 supplemented with 50μg/ml thymidine (thy) for overnight adaptation in culture tubes at 30°C. The next day (henceforth referred to as day 0; day 1 is the end of the first day of continuous culture), the overnight culture was used to inoculate three turbidostat tubes containing 17ml of M9 supplemented with 50 thy. The starting optical density was approximately 0.005. Each culture grew unperturbed until it reached an OD_600_ of 0.15, at which point it was diluted with 2.4 ml of fresh media. These cycles of growth and dilution persisted for a period of ~22 hours, at which point the run was paused to make glycerol stocks and replenish media. Culture vials for each following day of evolution were filled with fresh media and inoculated using 300μl from the previous culture.

### Growth Rate Measurements for DHFR and TYMS point mutants

All relative growth rate measurements were performed in the *E. coli* folate auxotroph strain ER2566 *ΔfolA ΔthyA* [60]. DHFR (*folA*) and TYMS (*thyA*) are provided on the plasmid pACYC-Duet1 (in MCS1 and MCS2, respectively) and are each under control of a T7 promoter. For these experiments, we use leaky expression (no IPTG induction). Each mutant plasmid (20 in total) is marked with a genetic barcode in a non-coding region between the two genes. Plasmids were transformed into the auxotroph strain, and each mutant was grown overnight in separate LB +30μg/ml Cam +50μg/ml thy cultures. Then, cultures were washed 2x in M9 media supplemented with 0.4% glucose and 0.2% amicase and 30μg/ml Cam, and adapted overnight at 30°C. All mutants were mixed in equal ratios based on OD_600_ and inoculated at a starting OD_600_ = 0.1 in the turbidostat. Growth rates were measured under two conditions: 5 thy and 50 thy, with three replicates each. The turbidostat clamps the culture to a fixed OD_600_ = 0.15 by adding fresh dilutions of media. Every 2 hours over the course of 12 hours a 1ml sample was removed, pelleted and frozen for next-generation sequencing. Amplicons containing the barcoded region with appropriate sequencing adaptors (350 basepairs in total size) were generated by two sequential rounds of PCR with Q5 polymerase. The barcoded region was sequenced with a single-end MiSeq run using a v2 50 cycle kit (Illumina). We obtained 14,348,937 reads. Data analysis was performed using a series of custom python scripts to count barcodes, and MATLAB to fit relative growth rates.

### Constructing DHFR/TYMS mutants in a clean genetic background

We followed the protocol for scarless genome integration using the modified λ-red system developed by Tas et al. [61]. In this method, a tetracycline (Tet) resistance cassette (“landing pad”) is first integrated at the site targeted for mutagenesis. Then, the landing pad is excised by the endonuclease I-SceI, and replaced with the desired mutation by *λ*-red mediated recombination. NiCl2 is used to counterselect against cells that retain the tetracycline cassette. Tas et al. provides a detailed protocol; here we give the specifics necessary for our experiments. For the *λ*-red machinery, we transformed the plasmid pTKRED (Genbank accession number GU327533) into electrocompetent *E. coli* MG1655 with a genomic *egfp/cat* resistance cassette (the forward evolution founder strain). For the Δ25-26 TYMS mutation, we introduced the *tetA* landing pad between genome positions 2,964,900 and 2,965,201 (genome NC000913) corresponding to the N-terminus of the *thyA* gene. For the DHFR mutations (L28R, W30R, and P21L), the landing pad was recombined between genome positions 49,684 and 49,990 (genome NC000913). In order to replace the Tet cassette, cells were induced with 2mM IPTG and 0.4% arabinose, and then transformed with 100ng of dsDNA PCR product containing the mutation of interest (with appropriate homology arms). This reaction experienced 3 days of outgrowth at 30°C in rich defined media (RDM, Teknova) with glucose substituted for 0.5% v/v glycerol. The media was supplemented with 6 mM or 4mM NiCl2 for counterselection against *tetA* at the *thyA* locus or *folA* locus respectively. The outgrowth culture was streaked onto agar plates and screened daily for the mutant of interest using LB supplemented with 50 μg/ml thy, 30 μg/ml spectinomycin, and +/- 5-10 μg/ml Tet. All mutations were confirmed by Sanger sequencing of the complete *folA* and *thyA* open reading frame; for *folA* the promoter region was also sequenced.

### LC-MS Metabolite Measurements

Cells were cultured in M9 0.2% glucose media containing 0.1% amicase, 50 ug/ml thy, and 30 ug/ml Cam at 30°C for metabolite analysis. In mid-log phase at OD_600_ ~0.2, *E. coli* culture (3 ml for nucleotide measurement and 7 ml for folate measurement) was filtered on a nylon membrane (0.2 μm), and the residual medium was quickly washed away by filtering warm saline solution (200 mM NaCl at 30’C) over the membrane loaded with cells to exclude non-desirable extracellular metabolites from LC-MS analysis. The membrane was immediately transferred to a 6 cm Petri dish containing 1 ml cold extraction solvent (−20°C 40:40:20 methanol/acetonitrile/water; for folate stability, 2.5 mM sodium ascorbate and 25 mM ammonium acetate in folate extraction solvent [45]) to quench metabolism. After washing the membrane, the cell extract solution was transferred to a microcentrifuge tube and centrifuged at 13000 rcf for 10 min. The supernatant was transferred to a new microcentrifuge tube. Folate samples were prepared with an additional extraction: the pellet was resuspended in the cold extraction solvent and sonicated for 10 min in an ice bath. After the second extraction and centrifugation, the supernatant was combined with the initial supernatant. The metabolite extracts were dried under nitrogen flow and reconstituted in HPLC-grade water for LC-MS analysis. Metabolites were measured using stand-alone orbitrap mass spectrometers (ThermoFisher Exactive and Q-Exactive) operating in negative ion mode with reverse-phase liquid chromatography [62]. Exactive chromatographic separation was achieved on a Synergy Hydro-RP column (100 mm×2 mm, 2.5 μm particle size, Phenomenex) with a flow rate of 200 μL/min. Solvent A was 97:3 H2O/MeOH with 10 mM tributylamine and 15 mM acetic acid; solvent B was methanol. The gradient was 0 min, 5% B; 5 min, 5% B; 7 min, 20% B; 17 min, 95% B; 20 min, 100% B; 24 min, 5% B; 30 min, 5% B. Q-Exactive chromatographic separation was achieved on an Poroshell 120 Bonus-RP column (150 x 2.1 mm, 2.7 μm particle size, Agilent) with a flow rate of 200 μl/min. Solvent A is 10mM ammonium acetate + 0.1% acetic acid in 98:2 water:acetonitrile and solvent B is acetonitrile. The gradient was 0 min, 2% B; 4 min, 0% B; 6 min, 30% B; 11 min, 100% B; 15 min, 100% B; 16 min, 2% B; 20 min, 2% B. LC-MS data were analyzed using the MAVEN software package [63].

## Supporting information

Supplemental Information - compiled

## Acknowledgements

We thank members of the Reynolds lab for review of the manuscript, E. Toprak for extensive advice on morbidostat construction and operation, S. Benkovic for the ER2566 *ΔfolA ΔthyA* strain, D. Bikard for the *E. coli* MG1655 strain containing dCas9, T. Kuhlman for molecular biology reagents used in genome editing, S. Thompson for the *folA* supplementation mix recipe, and T. Bergmiller for the GFP/Chloramphenicol resistance marker incorporated into our founder strain.

## Supplemental Legends

**Figure S1 Predicted interactions in folate metabolism from STRINGdb.** All data were obtained from the file COG.links.v10.5.txt, as downloaded from STRINGdb. **A**, The combined interaction, or confidence score between genes in central folate metabolism. The confidence score is benchmarked to approximate the probability that two enzymes share a pathway in the KEGG database. **B**, The neighborhood component of the interaction score, which captures genome context and chromosomal proximity. **C**, The co-occurrence component of the interaction score, based on correlations in COG presence/absence across species.

**Figure S2 Methods and validation for CRISPRi inhibition of expression. A**, Schematic of the vector containing sgRNAs for CRISPRi gene knockdowns. Each plasmid in the library contains two sgRNAs. For single gene knockdowns, an sgRNA that targets a gene is paired with an sgRNA lacking a targeting region (none). **B**, A representative sampling of frequency data and growth rate fits for the CRISPRi NGS-fit assay. Frequency is computed relative to the none:none sgRNA pair. **C**, The correlation between CRISPRi effects and gene knockout effects on growth suggests CRISPRi is working as expected. CRISPRi measurements are normalized to wildtype (none:none sgRNA), and were performed in M9 minimal media with glucose. Knockout growth data (x-axis) are published numbers associated with the KEIO collection (Baba et al, 2006 Mol Syst Biol), and report the normalized increase in OD_600_ over 24 hours of growth in MOPS minimal media with 0.4% glucose. **D**, RT-qPCR mRNA quantitation of CRISPRi effects on target gene mRNA abundance. All qPCR reactions were performed in technical triplicate except glyA, which was performed in technical duplicate. The ΔΔC_T_ method was used to quantity changes in gene expression. Standard error of the mean associated with each average C_T_ value was propagated using a first order Taylor expansion to produce the error bars associated with each gene knockdown.

**Figure S3 Evolution of trimethoprim (TMP) resistance in MG1655 cells using the morbidostat. A**, Schematic of a continuous culture tube. Dilutions were made through inlet tubes labeled ‘A’, ‘B’, and ‘C’. A constant volume of 15 ml was maintained by aspirating extra medium through the waste line after mixing. **B**, Control strategy for the addition of trimethoprim. Once the culture exceeded an OD_600_ of 0.06 periodic dilutions of 3 ml were made every 20 minutes. For OD_600_ between 0.06 and 0.15, media ‘A,’ containing no TMP, was added. While the OD_600_ was above a threshold of 0.15, media ‘B’ was used to introduce TMP. Media ‘B’ was added periodically until the culture tube reached a TMP concentration greater than 60% of the stock, at which point media ‘C,’ containing 5-fold more TMP was added. If a culture’s growth rate diminished to below that of the dilution rate, then media ‘A’ would be used in the next dilution to incrementally reduce the TMP concentration. At the end of a given day, if media ‘C’ had been utilized, then all stock concentrations of TMP were increased by a factor of 5X to enable continued adaptation in the following day. **C**, A representative growth trajectory, color-coded by TMP concentration (day 7, 50 μg/ml thymidine). See Figure S4 for full growth trajectories over 13 days. **D**, The trajectory of estimated TMP resistance (as measured by the median TMP concentration on each day) versus number of generations for each experimental condition (5, 10 and 50 μg/ml thymidine).

**Figure S4 OD600 measurements and trimethoprim concentration over 13 days of forward evolution.** Each plot corresponds to a different thymidine concentration, denoted in the upper left hand corner. The x-axis displays the number of days in real time. Discontinuities at day 5 for all three conditions are the result of a technical interruption; cultures were restarted from the previous day’s glycerol stock. An enhanced view of the 50 μg/ml trajectory day 7 (boxed in a dashed line) is shown in Fig S3B.

**Figure S5 Thymidine dependence of the 30 evolved strains.** The y-axis denotes the positive integral of log(OD_600_) evaluated over 20 hours of growth (see Experimental Procedures). Strains evolved for trimethoprim resistance in the presence of thymidine become auxotrophs.

**Figure S6 Lack of thymidine dependence after growth in the absence of TMP.** Loss of function in TYMS due to selection with TMP. **A**, Ten colonies from day 6 of the morbidostat TMP selection (50 μg/ml thymidine condition). Replica plating on 0 and 50 μg/ml thymidine indicates that all ten strains are thymidine auxotrophs. **B**, Ten colonies from three replicate growths in 50 μg/ml thymidine without TMP selection (turbidostat). These cultures were grown until biofilm formation became prohibitive. Replica plating on 0 and 50 μg/ml thymidine indicates that all strains retain TYMS activity.

**Figure S7 Strategy for measuring DHFR/TYMS mutant growth rates. A**, Genetic barcoding scheme for deep sequencing. Each plasmid contains two barcodes uniquely encoding the identity of its corresponding *folA* and *thyA* genotypes. **B**, Genotype frequency over time as determined by barcode sequencing. The relative fitness of each allele pair is given by the linear slope m. See Fig S8 for all growth rate fits.

**Figure S8 Relative growth rate measurements for DHFR/TYMS mutants.** Points represent the normalized relative frequency (log scale) of DHFR mutants during turbidostat growth in either 5 or 50 μg/ml thymidine. The y-axis indicates the genetic background of TYMS: either WT or R166Q. Relative growth rate fits are shown by the solid lines.

**Figure S9 The relationship between intracellular folate species and doubling time.** The y-axis depicts the log2 abundance of each metabolite normalized to WT. For each folate species, the five glutamylation states are shown in different colors. Doubling times were measured in M9 minimal media supplemented with 50 μg/ml thymidine. Error bars denote standard error of the mean across triplicate measurements.

