## Supplemental Information - compiled for "A two-enzyme adaptive unit within bacterial folate metabolism"

**Contents**

I. **Supplementary Figures**
   A. Figure S1: Predicted interactions in folate metabolism from STRINGdb
   B. Figure S2: Methods and validation for CRISPRi inhibition of expression
   C. Figure S3: Evolution of trimethoprim (TMP) resistance in MG1655 cells using the morbidostat
   D. Figure S4: OD600 measurements and trimethoprim concentration over 13 days of forward evolution
   E. Figure S5: Thymidine dependence of the 30 evolved strains
   F. Figure S6: Lack of thymidine dependence after growth in the absence of TMP
   G. Figure S7: Strategy for measuring DHFR/TYMS mutant growth rates
   H. Figure S8: Relative growth rate measurements for DHFR/TYMS mutants
   I. Figure S9: The relationship between intracellular folate species and doubling time

II. **Supplementary Experimental Procedures**
   A. Analysis of synteny and co-occurrence

B. Extended methods for CRISPR interference (CRISPRi) and next-generation sequencing based measurements of relative growth rate

**III. Supplementary Tables**

A. Table S1: Enzymes in central folate metabolism
B. Table S2: Primers used for CRISPRi library construction and sequencing
C. Table S3: sgRNA sequences and homology locations
D. Table S4: CRISPRi and sgRNA plasmids
E. Table S5: qPCR primers for quantifying CRISPRi efficiency
F. Table S6: Trimethoprim resistance (IC50) for forward evolution strains
G. Table S7: Whole genome sequencing statistics
H. Table S8: Complete list of mutations during the forward evolution experiment
I. Table S9: Functional annotations for commonly mutated genes
J. Table S10: Modular protein pairs identified by gene synteny

Figure S1
Schober et al.

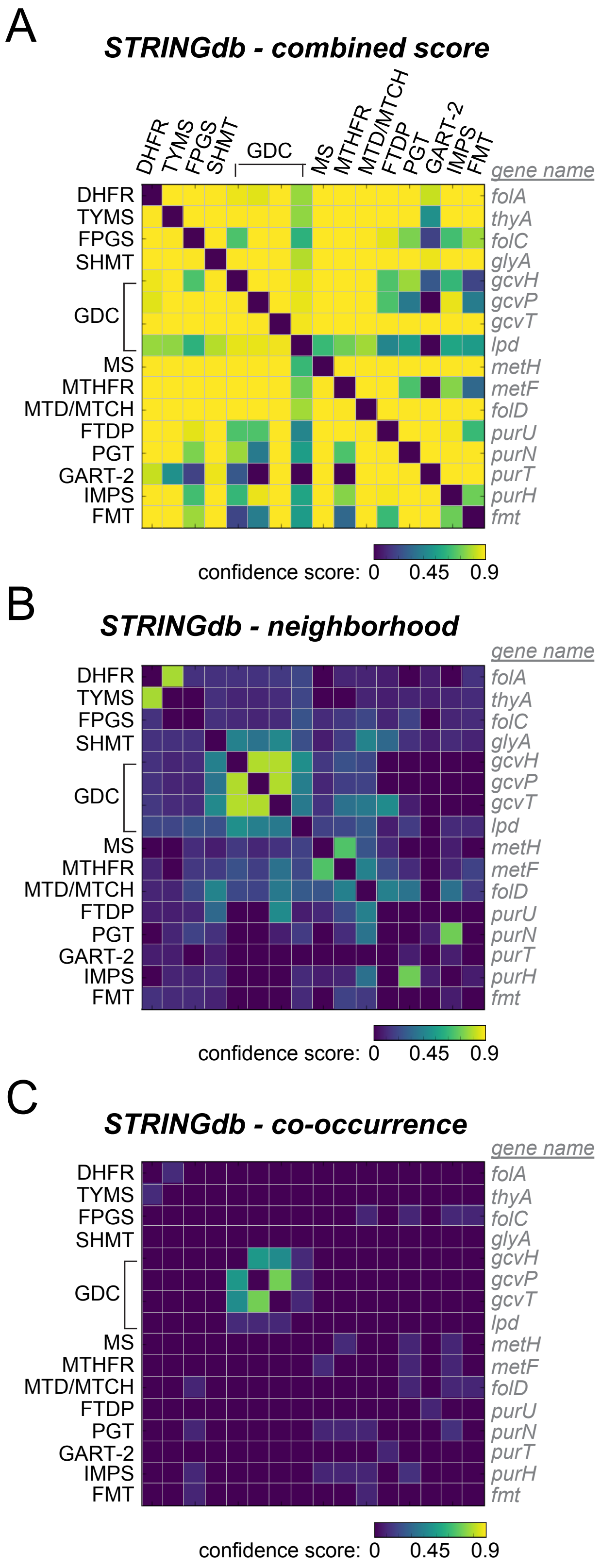

Figure S2
Schober et al.

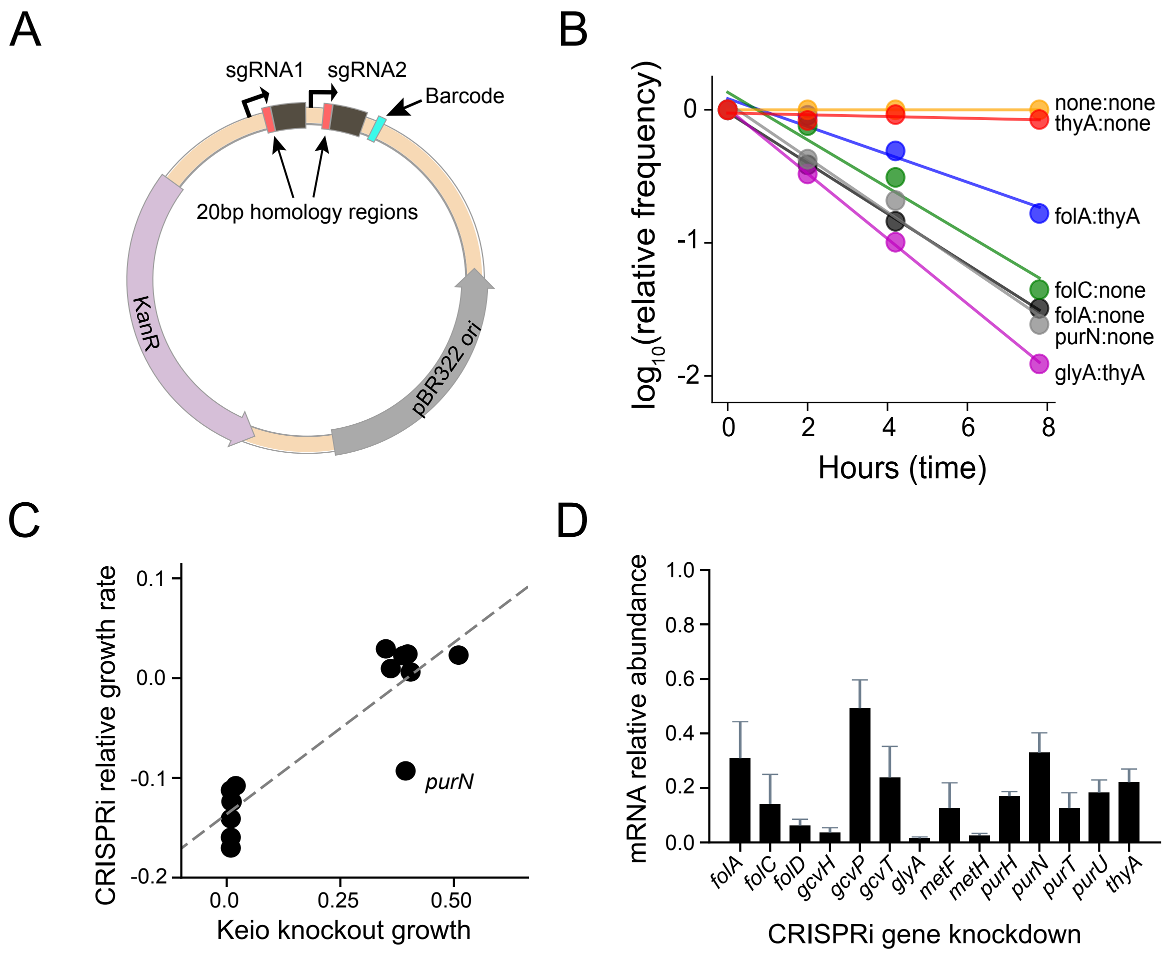

Figure S3
Schober et al.

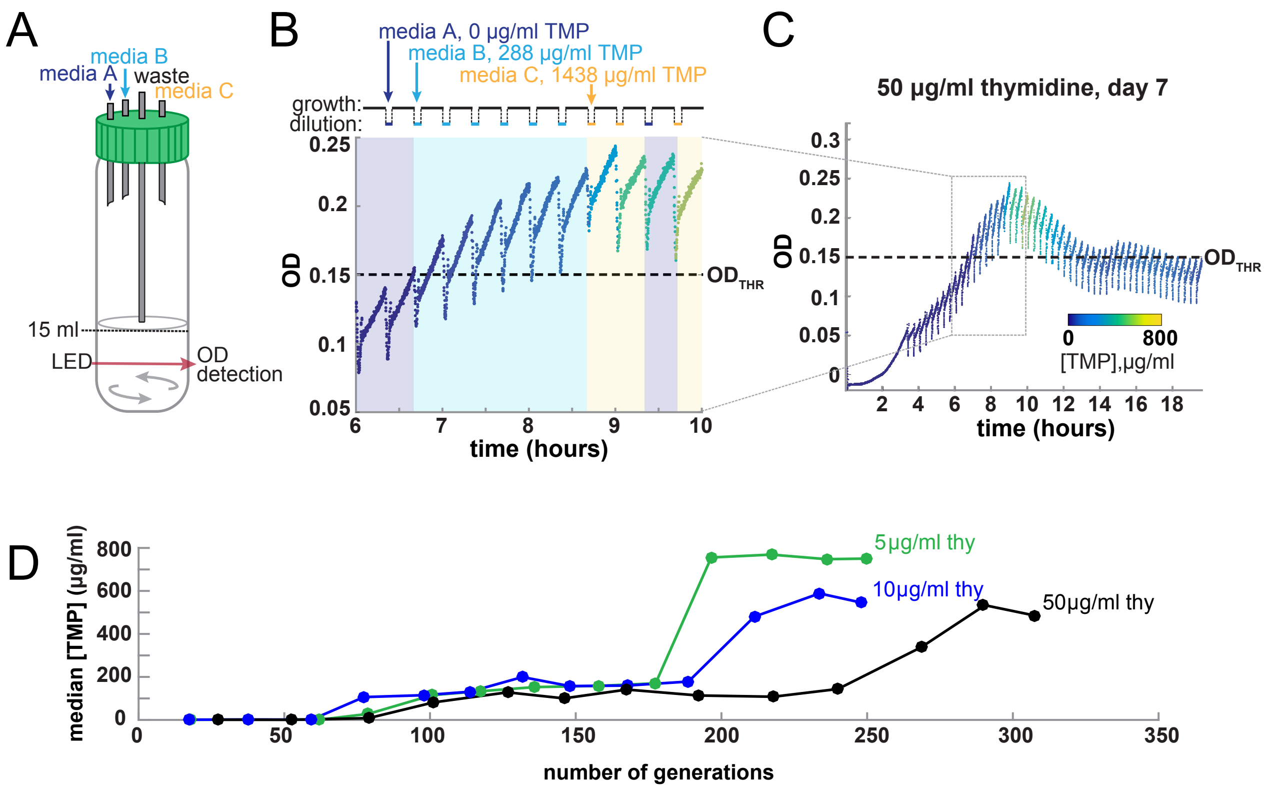

Figure S4
Schober et al.

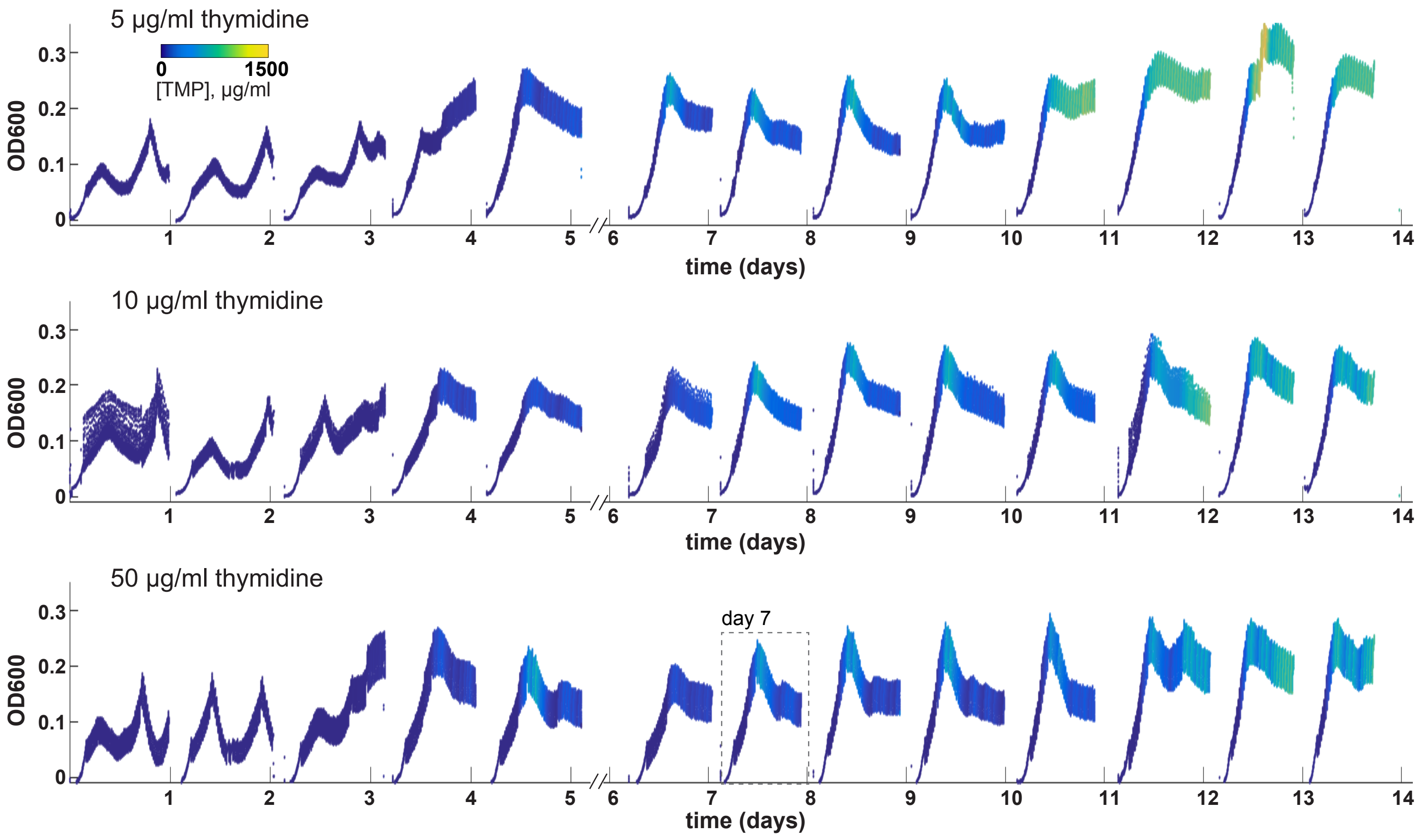

Figure S5
Schober et al.

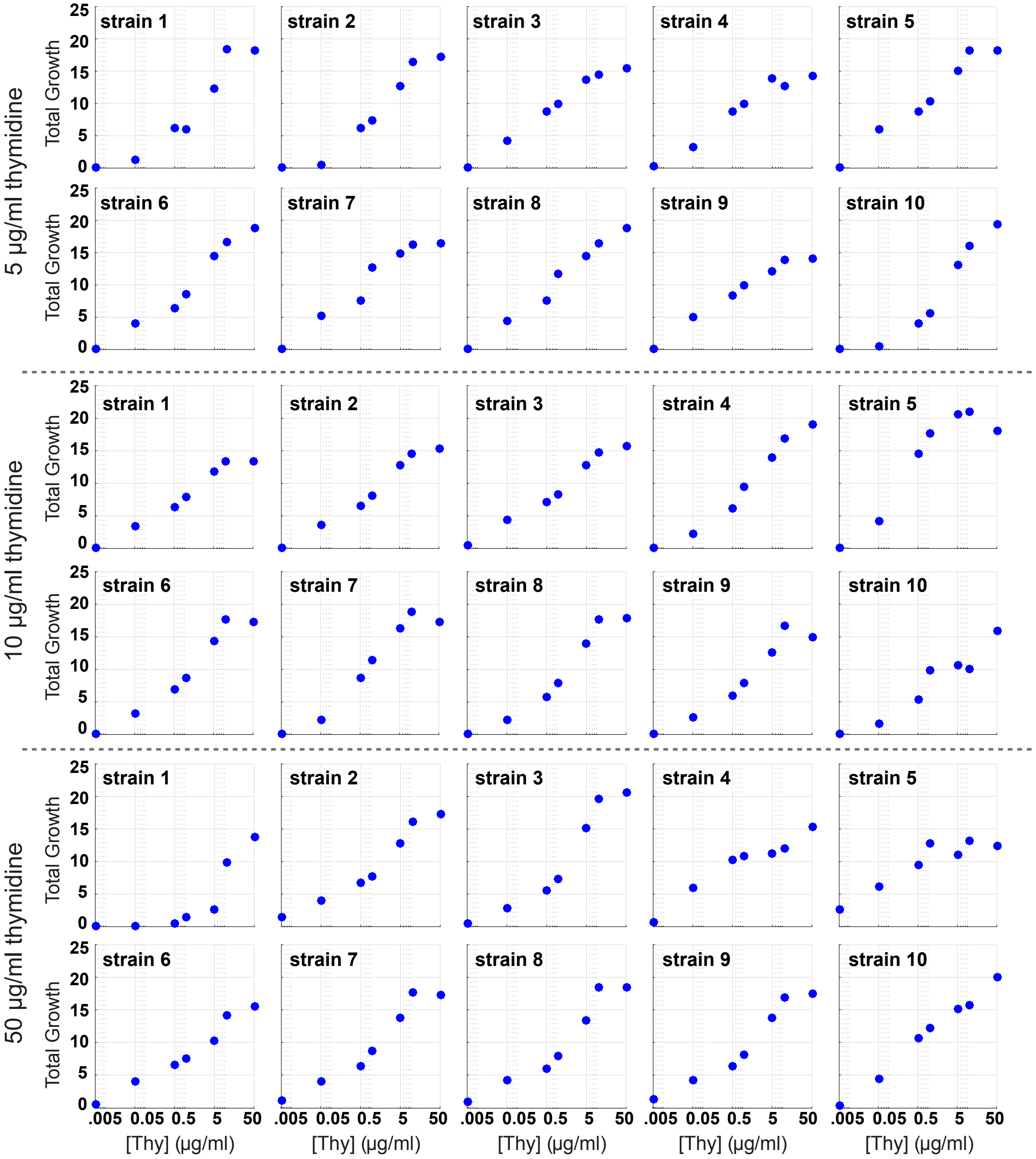

Figure S6
Schober et al.

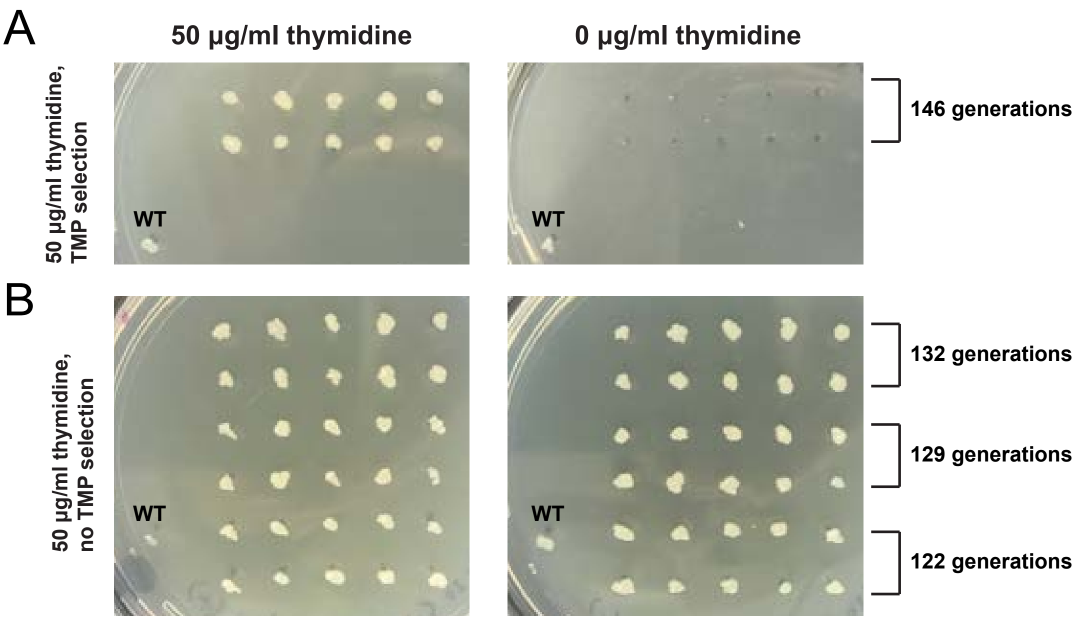

Figure S7
Schober et al.

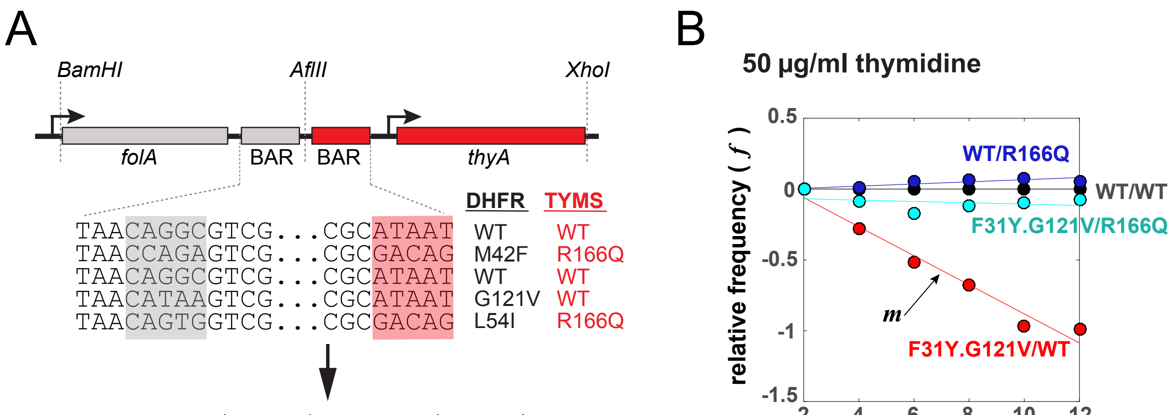


$$f(t) = log\left(\frac{N_{Mut}}{N_{WT}}\right)_t - log\left(\frac{N_{Mut}}{N_{WT}}\right)_{t=0}$$

Figure S8
Schober et al.

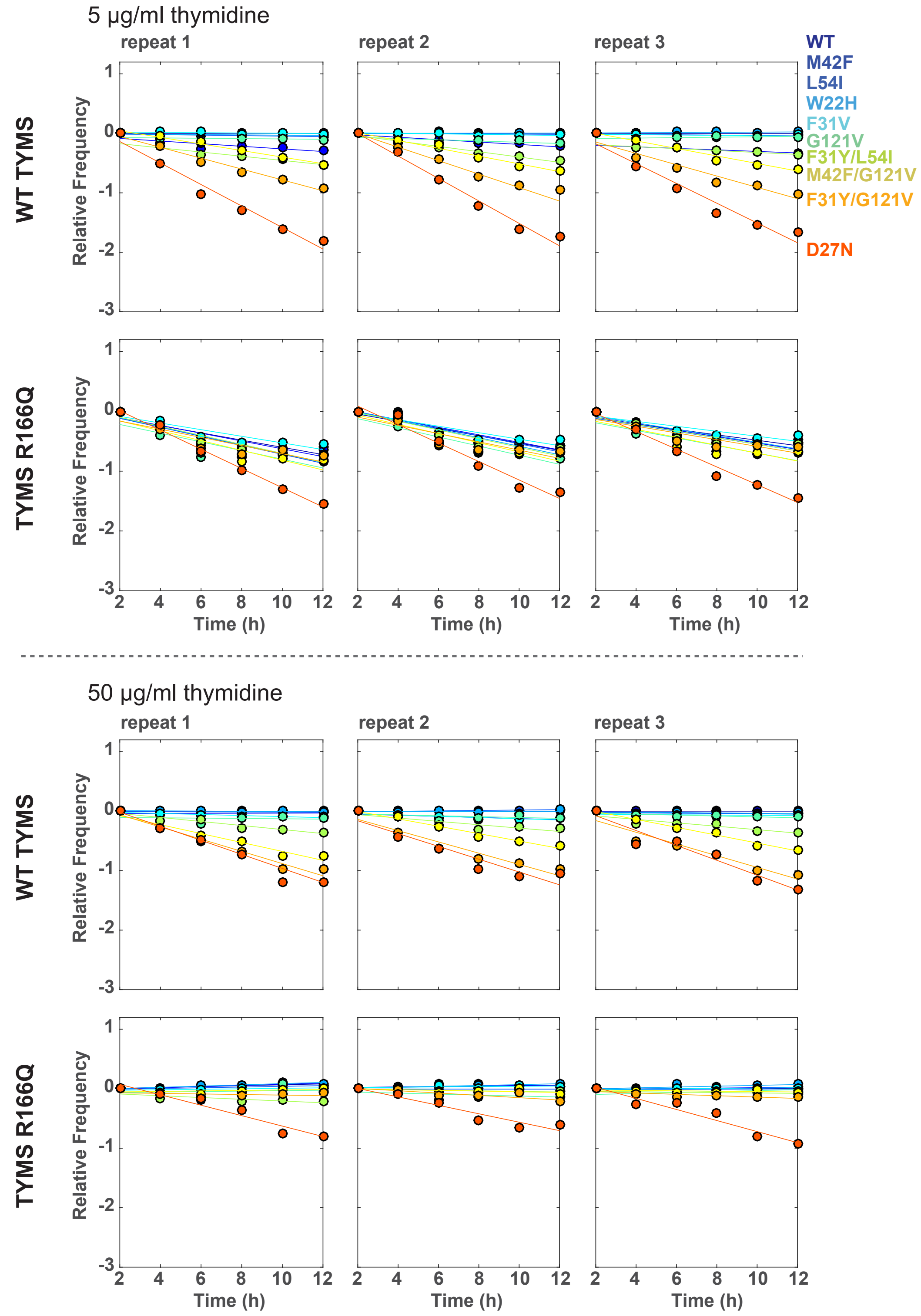

Figure S9
Schober et al.

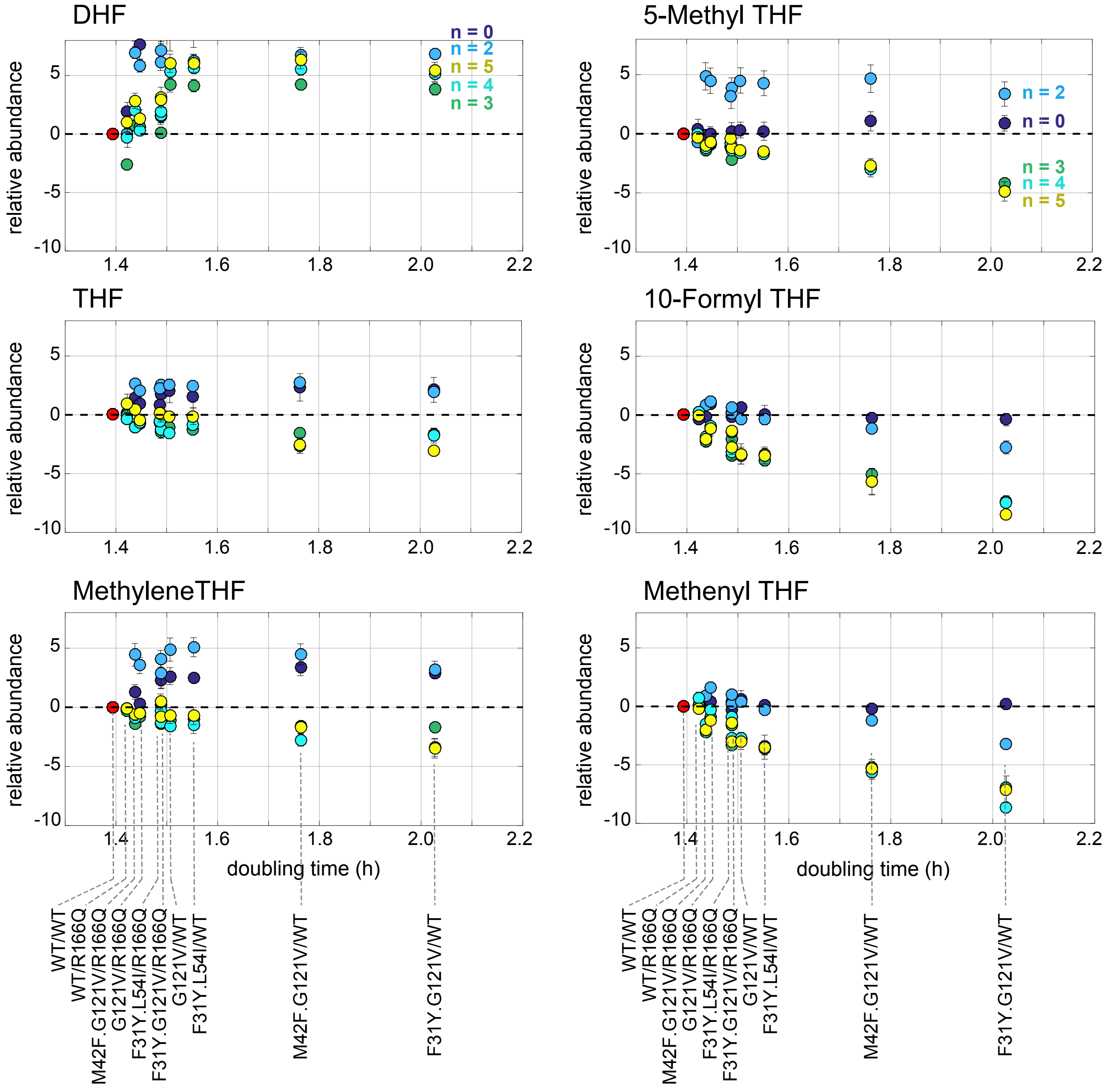

### Calculations of Synteny and Co-occurrence

#### A. Starting dataset

Calculations of both synteny and co-occurrence require a collection of genomes where individual genes are assigned into orthology classes. The Clusters of Orthologous Groups of proteins (COGs) defined by Koonin and colleagues provide one well-established set of ortholog annotations (Galperin et al., 2015). The results presented here use all complete and COG-annotated bacterial genomes available in the NCBI database as of March 2015 (1445 genomes and 4764 COGs, this dataset is also used in Junier and Rivoire, 2016). A genome may contain more than one gene in the same COG, but for clarity, we start by presenting the calculations assuming that every orthology class maps to at most one gene in each genome.

#### B. Counting pairs in co-occurrence

We begin by counting the number of genomes where orthology classes $i$ and $j$ co-occur. As previously published (Junier and Rivoire, 2016), we correct for the uneven phylogenetic distribution of sequenced genomes (strains) by introducing genome weights. To this end, we compute a distance between each pair of strains, based on the sequence similarity of a few conserved genes ($\delta_{gh} = 1 - S_{gh}$, where $S_{gh}$ is the average sequence similarity). The weight $w_s$ of strain $s$ is then defined as $1/n_s$ where $n_s$ is the number of strains within a given distance $\delta$ of $s$. Varying $\delta$ can provide information at different "phylogenetic depths" (Junier and Rivoire, 2013) but here we fix $\delta = 0.3$, our results being generally invariant to this value.

The effective number of strains where orthology classes $i$ and $j$ co-occur is formally given by

$$M_{ij} = \sum_s w_s \mathbb{1}[i \cap s \neq \emptyset]\mathbb{1}[j \cap s \neq \emptyset] \tag{1}$$

where the sum is over the strains $s$ and where $\mathbb{1}[X]$ is a generic indicator function with $\mathbb{1}[X] = 1$ if and only if $X$ is true. Hence, $\mathbb{1}[i \cap s \neq \emptyset] = 1$ if $i$ is represented in strain $s$ and 0 otherwise.

#### C. Defining gene proximity

We measure the distance $d(i, j)$ between the midpoint of two genes $i$ and $j$ in base pairs (and set $d(i, j) = \infty$ if they are on different chromosomes). Given a circular chromosome of length $L$, the greatest possible distance between genes is $L/2$ (on opposite sides of the circle). Thus, given a null model in which genes are randomly distributed along the chromosome, the probability of finding the gene pair within a genomic proximity $d^*$ is just the normalized value $p^* = d^*/(L/2)$.

#### D. Counting pairs in synteny

The value $p^*$ provides a measure of signifcance for finding two genes at a distance $d^*$ in one genome. However, we are interested in the conservation of proximity across many species. To begin, we count the effective number of strains in which $i$ and $j$ are within a given distance $d^*$.

$$X_{ij} = \sum_s w_s \mathbb{1}[d(i, j) < d^*], \tag{2}$$

However, because $p^* = (2d)/L$, the probability of finding two genes within distance $d^*$ depends on the chromosome length $L$, which varies between strains. In order for the probability of observing a positive event under the null model to be common for all strains, we instead consider the normalized distance and compute:

$$X_{ij} = \sum_s w_s \mathbb{1}[2d(i, j)/L_s < p^*]. \tag{3}$$

For strains that contain multiple chromosomes, we take for $L_s$ the sum of the lengths of its different chromosomes. This corresponds to a null model where the genes are randomly shuffled within and between chromosomes (or, up to boundary effects, to concatenating all the chromosomes into a single one). We take $p^* = 0.02$, corresponding to $d = 50$ kb in the context of a chromosome of length 5 Mb. This cutoff is chosen to represent a length scale longer than those typical for gene coexpression and synteny, so that the choice of cutoff does not determine the results. Further, the results are robust with respect to the choice of $p^*$.

Finally, to account for the possibility that a single strain may contain multiple pairs of genes in two given orthology classes $ij$, we correct Eq. (3) by averaging over all these pairs:

$$X_{ij} = \sum_s w_s \frac{1}{|i \cap s||j \cap s|} \sum_{g_i \in i \cap s, g_j \in j \cap s} \mathbb{1}[2d(g_i, g_j)/L_s < p^*], \tag{4}$$

where $i \cap s$ is as before the set of genes in orthology class $i$ and in strain $s$ and $|i \cap s|$ the size of this set. This formula is simpler than the one used in (Junier and Rivoire, 2016) but leads to similar results.

#### E. Measuring significance

Now that we have counted the number of genomes in which $i$ and $j$ are proximal, we can assess the significance of this result. In a standard statistics "coin toss" problem, one computes the significance of obtaining $X$ "tails" out of $M$ "flips" (given a probability of tails $p^* = 0.5$) using the binomial distribution. Here, we compute the significance of finding a pair of genes in proximity $X_{ij}$ times out of $M_{ij}$ genomes (given a probability of $p^* = 0.02$) using the same approach:

$$\pi_{ij} = \sum_{K \geq X_{ij}} \binom{M_{ij}}{K} (p^*)^K (1-p^*)^{M'_{ij}-K} = I(X_{ij}, M_{ij} - X_{ij} + 1, p^*) \tag{5}$$

where $I(a, b, x)$ is the regularized incomplete beta function.

This relatively naive null model (which assumes a uniform distribution of genes along the chromosome, and treats weighted genomes as independent trials) provides a good description of the data for the majority of orthology class pairs - indicating that most gene pairs have no significant conservation of chromosomal proximity (Junier and Rivoire, 2016). A subset of pairs nevertheless deviate from the statistical expectations of the null model; these are the syntenic pairs of interest.

Finally, analysis of any large dataset inevitably leads to spurious false positives that simply occur by random chance. To account for this, we apply the Bonferroni principle - we set here a threshold of significance to $\pi^* = 2/N(N-1) \simeq 10^{-7}$ where $N = 4764$ is the number of orthology classes defined by COGs. That is, we choose a cutoff such that we should not find any significantly syntenic gene pair "by random" among all $10^7$ possible gene pairs. This criterion is very stringent, and may be relaxed to set instead a false discovery rate (Junier and Rivoire, 2016).

#### F. Degree of synteny

The p-values $\pi_{ij}$ depend on the number of genomes in the dataset. It is more meaningful to define a measure of conservation that depends only on rescaled variables, here the frequencies $f_{ij} = X_{ij}/M_{ij}$. For these frequencies to be meaningful, we need, however, to restrict to cases where the number $M_{ij}$ of genomes where genes $i$ and $j$ co-occur is large. Here, we restrict to pairs of COGs with $M_{ij} \geq 100$. A degree of synteny is then given by the relative entropy:

$$D(f_{ij} \| p^*) = f_{ij} \ln \frac{f_{ij}}{p^*} + (1 - f_{ij}) \ln \frac{1 - f_{ij}}{1 - p^*}. \tag{6}$$

In the limit of large $M_{ij}$, $e^{-M_{ij} D_{ij}}$ approximates the first term of the sum in Eq. (5) and therefore $M_{ij} D_{ij}$ correlates with $-\ln \pi_{ij}$. The maximal value of $D_{ij}$ is set by $p^*$: as $p^* = 0.02$ corresponds to $-\ln p^* \simeq 4$, the range of values for $D_{ij}$ is thus $[0, 4]$. Finally, since $M_{ij} \geq 10^2$ and $\pi^* = 10^{-7}$, any value of $D_{ij}$ larger than $D^* = -(\ln 10^{-7})/10^2 \simeq 0.025$ reports significant synteny.

##### G. Degree of co-occurrence

Following the same logic, we can simlarly define a measure for co-occurrence. We compute the probability of finding a given gene pair $i, j$ together (in co-occurrence) $X_{ij}$ or more times in a set of $M$ genomes, given the null model that the two genes are independent ($f_{ij} = f_i f_j$). Again beginning from the binomial distribution function, and taking Stirling's approximation, one obtains the mutual information as a measure of co-occurrence between genes:

$$D(f_{ij}|f_i f_j) = f_{ij} \ln \frac{f_{ij}}{f_i f_j} + (1 - f_{ij}) \ln \frac{1 - f_{ij}}{1 - f_i f_j} \tag{7}$$

##### H. Application to *E. coli*

To analyze synteny and co-occurrence relationships relevant to *E. coli*, we keep only the COGs $i$ that are represented in its genome, and analyze COG pairs for which $M_{ij} \geq 100$ (2095 COGs in total). In Fig. 6, we plot for each pair $ij$ of these COGs their degree of synteny (or co-occurrence) $D_{ij}$ (x-axis) against their maximal degree of synteny (or co-occurrence) with any other COG $\max_{k \neq i,j}(D_{ik}, D_{jk})$ (y-axis). In this figure, we annotate physical interactions using the STRING "actions" database for *E. coli* (511145.protein.actions.v10.5.txt - this file reports the subset of STRING interactions which correspond to physical binding) taking a threshold of 700 ("high confidence") and the largest score when multiple paralogs are present. To annotate genes within the same pathway and enzyme pairs sharing an intermediate, we use KEGG pathways/reactions. Because a number of the KEGG pathways are extremely general, we removed the four most inclusive pathways from consideration of "shared pathway" status: "Metabolic Pathways", "Biosynthesis of secondary metabolites", "Microbial metabolism in diverse environments", "Biosynthesis of antibiotics". For the annotation of shared metabolites, we also removed all KEGG compounds that occur in 50 reactions or more (H2O, ATP, ADP, Orthophosphate, Diphosphate, H+, NAD+, NADH, NADPH, NADP+, CO2, NH3, AMP).

**Extended Methods for CRISPR interference (CRISPRi) and next generation sequencing based measurements of relative growth rate.** We used a combination of Clustered Regularly Interspaced Short Palindromic Repeats interference (CRISPRi) and Next-Generation Sequencing (NGS) to measure the effects of gene knockdown on bacterial growth; we refer to this assay as NGS-Fit for brevity. We describe the methods for this approach in two parts: sgRNA library construction, and the fitness assay.

**sgRNA Library Construction.** CRISPRi sgRNAs were designed to target 14 genes in *E. coli* central metabolism (*folA*, *thyA*, *folC*, *glyA*, *gcvH*, *gcvP*, *gcvT*, *metH*, *metF, folD*, *purU*, *purN, purT*, and *purH)*. All the genes except for *gcvP*, *gcvT*, and *gcvH* are in different operons, ensuring separate targeting. For each gene, we selected a 20bp homology region (preceding a PAM site) on the non-template strand of the 5' end of the gene for targeting. We used BLAST to confirm that each guide lacks significant homology to any other locus in the genome. See Table S3 for sequences and PAM sites. The individual sgRNAs were cloned into a modification of the pCRISPR plasmid (Addgene #42875). The modified pCRISPR plasmid was made by removing the original sgRNA sequences by restriction digest and replacing them with a small DNA sequence containing two BsaI sites for golden gate cloning (creating pAM-111, Table S4). The sgRNA (and corresponding promotor) from the pgRNA vector (Addgene #44251) was then inserted into pAM-111 using golden gate cloning (creating pAM-112, Table S4). From this point forward, making a new sgRNA was done by inserting a new homology region into pAM-112 using iPCR as described (Table S2, Hawkins et al, Methods Mol. Biology 2015;1311:349).

To achieve pairwise knockdowns, we used golden gate assembly to combine the sgRNA inserts for *folA* (or *thyA*) with all other genes in folate metabolism or with a negative control guide (*none*) which lacks a targeting homology region. The primers GG-sgRNA1-F/R or GG-sgRNA2-F/R were used to amplify the sgRNA inserts with BsaI sites from pAM-112 for the golden gate reaction (Table S2). The inserts were then ligated into three vectors (pAM-288, pAM-289, and pAM-290) using three separate golden gate reactions. pAM-288, 289, 290 are nearly identical to pAM-111 with two important differences. First, orthogonal primer sites were inserted into this vector flanking the golden site. These orthogonal primers allow specific amplification of plasmids that have the golden gate BsaI sites and do not amplify the pAM-112 parent plasmid which carries over from the sgRNA insert amplification. Second, pAM-288, 289, 290 each contain a unique 6N barcode (CTTTCA, ATCATG, GCATGG) directly downstream of the sgRNAs. Thus, following assembly, we obtain three small libraries of 39 unique knockdowns, each with a unique barcode. Because the sgRNAs can be cloned in either order in the plasmid, we end up with six total replicates of each gRNA pair in our full library (three barcodes in two orders). This allows us to calculate the error across internal replicates in a single experiment, and potentially exclude so-called "escapers". The final library was transformed into MG1655 *E. coli* with dCas9 in the chromosome (gift from Bikard lab) and glycerol stocked (gAM-350).

**Next-generation sequencing-based fitness assays (NGS-fit).** Immediately following transformation, the cells were grown over night at 37 °C shaking in a SOB LB mix containing 35 μg/mL Kanamycin, folA mix [38 μg/mL glycine, 75.5 μg/mL methionine, 1 μg/mL pantothenate, 20 μg/mL adenosine], and 50 μg/mL thymidine. These additives relieve selection pressure on genes targeted by CRISPRi during outgrowth in case of leaky dCas9 expression. In the morning, the culture was centrifuged and washed 2 times with M9 minimal media pH 6.5, 0.4% glucose, 35 µg/mL Kanamycin, 5 µg/mL thymidine, and 0.2% amicase. The cells were adapted to this media for 12 hours at 30 °C shaking. The cultures were then back diluted to OD600 0.05 and used to inoculate 15 ml cultures in our turbidostat at 30 °C. Optical density was clamped at an OD600 of 0.15. Turbidostat input media was M9 minimal media pH 6.5, 0.4% glucose, 35 µg/mL Kanamycin, 5 µg/mL thymidine, and 0.2% amicase.

The culture was further adapted in the turbidostat for 8 hours, then CRISPRi was induced by the addition of 50 ng/mL anhydrous tetracycline to the media (aTC). Culture samples (1 ml each) for NGS were taken 3 hours, 5 hours, 7 hours, and 11 hours after CRISPRi induction. These samples pelleted by centrifugation, decanted, and frozen at -20 °C. All samples were prepared for amplicon sequencing using two rounds of PCR to add standard Illumina TruSeq and i5/i7 sequencing primers. Each time point was assigned a unique i5/i7 combination for de-multiplexing after sequencing. PCR amplicons for each sample (time point) were quantified using a picogreen assay and then mixed in equal stoichiometric ratio. This mix was run on an agarose gel and the band corresponding to the expected size (~550 BP) was excised. The resulting amplicon mixture was sequenced on an Illumina miSeq using a 300 cycle V2 kit configured to read 175 base pairs (read 1) and 125 base pairs (read 2) to sequence the barcode and both sgRNA homology regions. MiSeq quality: reads passing Q30=96.4%, 818K/mm$^2$ cluster density, and 15,384,044 reads passing filters.

Data from the miSeq was processed using a custom data analysis script written in Python 2.7. This script counts how often each sgRNA pair and barcode combination were found at each time point. Count data is normalized to the none|none sgRNA pair at each time point (to control for mixing variation), and to the frequency (*f*) at the first time point:

$$f(t) = log_{10}\left(\frac{n_{sg}}{n_{none}}\right)_t - log_{10}\left(\frac{n_{sg}}{n_{none}}\right)_{t=0}$$

Where n is the number of NGS counts of a given sgRNA (sg) or the none sgRNA (none) at a time t. We then conduct a linear fit of the relative frequency vs. time to determine a relative growth rate for each sgRNA combination.

**q-PCR methods for CRISPRi sgRNA verification.** CRISPRi gene knockdowns were induced in MG1655 *E. coli* growing in M9 minimal media (0.4% glucose, 35 µg/ml Kanamycin) at 37 °C with 50 ng/µl ATc for four hours prior to RNA extraction. Total RNA extractions were performed using the Purezol reagent (Bio-Rad, 7326890) following manual instructions. rt-qPCR was performed using the Luna Universal One-Step RT-qPCR kit (NEB, cat #E3005S) and a CFX384 Real-Time System, both following manual instructions. The *hcaT* gene was used as the house keeping control. Every reaction was performed in technical triplicate except the *glyA* knockdown measurement, which was performed in duplicate. Melt curves were performed for all qPCR primer pairs to confirm single product amplification. However, if in later experiments one of the technical replicates had a multi-modal melt curve, it was removed. Relative mRNA expression was calculated using the $2^{-\Delta\Delta Ct}$ method, and standard error of the mean from technical replicates was propagated using a first order Taylor Series expansion.

| Abbreviation | Name | E. coli gene | Uniprot ID | COG |
|---|---|---|---|---|
| **FPGS** | bifunctional dihydropteroate synthase/FPGS | *folC* | P08192 | COG0285H |
| **DHFR** | dihydrofolate reductase | *folA* | P0ABQ4 | COG0262H |
| **TYMS** | thymidylate synthase | *thyA* | P0A884 | COG0207F |
| **SHMT** | serine hydroxymethyltransferase | *glyA* | P0A825 | COG0112E |
| **GDC** | glycine cleavage system - protein H | *gcvH* | P0A6T9 | COG0509E |
| | glycine cleavage system - protein L | *lpdA* | P0A9P0 | COG1249C |
| | glycine cleavage system - protein P | *gcvP* | P33195 | COG1003E |
| | glycine cleavage system - protein T | *gcvT* | P27248 | COG0404E |
| **MS** | methionine synthase | *metH* | P13009 | COG1410E |
| **MTHFR** | 5,10-methylenetetrahydrofolate reductase | *metF* | P0AEZ1 | COG0685E |
| **MTD** | methylenetetrahydrofolate dehydrogenase (bifunctional) | *folD* | P24186 | COG0190H |
| **MTCH** | methenyltetrahydrofolate cyclohydrolase (bifunctional) | *folD* | P24186 | COG0190H |
| **FTDP** | formyltetrahydrofolate deformylase | *purU* | P37051 | COG0788F |
| **PGT** | phosphoribosylglycinamide formyltransferase | *purN* | P08179 | COG0299F |

Table S1 **Enzymes in central folate metabolism**.

| Primer name | Sequence |
|---|---|
| **folA-sgRNA** | AGATCGGCAGGCAGGTTCCAGTTTTAGAGCTAGAAATAGCAAGTTAAAATAAGGC |
| **thyA-sgRNA** | AAAGCGTTCCGGTTCCGGTAGTTTTAGAGCTAGAAATAGCAAGTTAAAATAAGGC |
| **folC-sgRNA** | GCCAGAGGCGACGCGGCTTGGTTTTAGAGCTAGAAATAGCAAGTTAAAATAAGGC |
| **glyA-sgRNA** | CCACAGTTCGGCATCATAATGTTTTAGAGCTAGAAATAGCAAGTTAAAATAAGGC |
| **gcvH-sgRNA** | TTGCTGTATTTCAGTTCTGCGTTTTAGAGCTAGAAATAGCAAGTTAAAATAAGGC |
| **gcvP-sgRNA** | GCGCGGCGTCCGGTCCGATAGTTTTAGAGCTAGAAATAGCAAGTTAAAATAAGGC |
| **gcvT-sgRNA** | AGCGTGTGTTGTTCGTACAAGTTTTAGAGCTAGAAATAGCAAGTTAAAATAAGGC |
| **metH-sgRNA** | AAAGCGTTCACCACGAAAATGTTTTAGAGCTAGAAATAGCAAGTTAAAATAAGGC |
| **metF-sgRNA** | ATTCAGGGCATCCCGCTGGCGTTTTAGAGCTAGAAATAGCAAGTTAAAATAAGGC |
| **folD-sgRNA** | GTTACTACCCACCAGCACAAGTTTTAGAGCTAGAAATAGCAAGTTAAAATAAGGC |
| **purU-sgRNA** | TAGTACGCAGAACTTTACGTGTTTTAGAGCTAGAAATAGCAAGTTAAAATAAGGC |
| **purN-sgRNA** | CTGTAAATTACTTCCGTTGCGTTTTAGAGCTAGAAATAGCAAGTTAAAATAAGGC |
| **purH-sgRNA** | CTGAGCAGAGCTCGGCGGACGTTTTAGAGCTAGAAATAGCAAGTTAAAATAAGGC |
| **folM-sgRNA** | AATTAATATTGGCAAGGGCTGTTTTAGAGCTAGAAATAGCAAGTTAAAATAAGGC |
| **purT-sgRNA** | AACATCACGCGAGTTGCTGCGTTTTAGAGCTAGAAATAGCAAGTTAAAATAAGGC |
| **negC-sgRNA** | GTTTTAGAGCTAGAAATAGCAAGTTAAAATAAGGC |
| **Universal R-sgRNA** | ACTAGTATTATACCTAGGACTGAGCTAGC |
| **prAM-328 (TruSeq R)** | CACTCTTTCCCTACACGACGCTCTTCCGATCTNNNNCCATTATTAGTACAGCGAGGCAAC |
| **prAM-331 (TruSeq F)** | TGACTGGAGTTCAGACGTGTGCTCTTCCGATCTNNNNGTACGACTCCATGTTATGTATGCC |
| **GG-sgRNA1-F** | GTACTGGGTCTCTAGGTTTGACAGCTAGCTCAGTCCTAG |
| **GG-sgRNA1-R** | GTACTGGGTCTCTGGTACTTCAAAAAAAGCACCGACTCG |
| **GG-sgRNA2-F** | GTACTGGGTCTCTTACCTTGACAGCTAGCTCAGTCCTAG |
| **GG-sgRNA2-R** | GTACTGGGTCTCTCCACCTTCAAAAAAAGCACCGACTCG |

Table S2 **Primers used for CRISPRi library construction and sequencing.** Sequences are specified in the 5' to 3' direction.

| gene_name | BP_Region | PAM_Found | PAM_BP_location |
|---|---|---|---|
| **purN** | CTGTAAATTACTTCCGTTGC | CCG | 23 |
| **glyA** | CCACAGTTCGGCATCATAAT | CCG | 26 |
| **thyA** | AAAGCGTTCCGGTTCCGGTA | CCG | 60 |
| **gcvP** | GCGCGGCGTCCGGTCCGATA | CCA | 51 |
| **gcvH** | TTGCTGTATTTCAGTTCTGC | CCA | 13 |
| **gcvT** | AGCGTGTGTTGTTCGTACAA | CCT | 16 |
| **metF** | ATTCAGGGCATCCCGCTGGC | CCA | 17 |
| **purU** | TAGTACGCAGAACTTTACGT | CCA | 12 |
| **folM** | AATTAATATTGGCAAGGGCT | CCC | 11 |
| **purT** | AACATCACGCGAGTTGCTGC | CCG | 28 |
| **folC** | GCCAGAGGCGACGCGGCTTG | CCT | 19 |
| **folA** | AGATCGGCAGGCAGGTTCCA | CCG | 61 |
| **purH** | CTGAGCAGAGCGCGGCGGAC | CCA | 16 |
| **folD** | GTTACTACCCACCAGCACAA | CCG | 110 |
| **metH** | AAAGCGTTCACCACGAAAAT | CCG | 104 |
| **negC** | no targeting region | | |

Table S3 **sgRNA sequences and homology locations.**

| Plasmid | Purpose | Genbank File |
|---|---|---|
| pAM-111 | modified pCRISPR plasmid with BsaI sites | pAM-111 |
| pAM-112 | pAM-111 with 1 sgRNA inserted by Golden Gate | pAM-112 |
| pAM-288/290 | Modified pAM-111 with new primer binding sites and unique 6 base pair barcodes | pAM-288 |
| pAM-350 | Example of 2 sgRNAs inserted into pAM-288 with golden gate cloning | pAM-350-ex |

Table S4 **CRISPRi and sgRNA plasmids.**

| Gene | F primer | R primer |
| --- | --- | --- |
| *folA* | TGGGAATCAATCGGTCGTCC | ACCGACTTCACCCACGTTAC |
| *folC* | TCCAGATTGTGAGCGAGTCG | TTCAACCAGGCCAGAGTTCC |
| *folD* | CGTCTCCCGCTCTTATGACC | TCCACGTCTTTGTCCGGATG |
| *gcvH* | AAAGAACACGAATGGCTGCG | GACGCCGCTTTTACCGATTC |
| *gcvP* | GGTATACCGCGTACACTCCG | GCAGAGGCCATATCCAGTCC |
| *gcvT* | TCCGCCTCGTTGTTAACTCC | ACGGTTTCATCCCTTCCACC |
| *glyA* | CACCTGACTCACGGTTCTCC | TTTCACGCATTTTCGCCCAG |
| *metH* | TGGAAGATGAAGTGGTGGGC | GGATTCAGCGTTTTCTCGGC |
| *metF* | ATGCTTCTGACCTGGTGACG | AGAACTGAGTAATCGCGCGG |
| *purH* | CGCCATCAAAGCCTTCGAAC | TCTCGCCGTAACGCATATCC |
| *purN* | TTACACACCCATCGTCAGGC | GCAAATACCGGGACTTTCGC |
| *purT* | TGCCACCGATATGCTGATCC | TAAGTGGAAGTGGGCAGCTG |
| *purU* | TGACAATCTGGACGAAGGCC | GCGCGCATCATATCTTCAGC |
| *thyA* | GCTCCTGTGACGTCTTCCTC | ATCACCCACTTCCAGATCGC |
| *hcaT* | CTTTTCCGCCGTTGTAGTGC | GGATGACTTCGCTACCCTGG |

**Table S5 qPCR primers for quantifying CRISPRi efficiency.** Sequences are specified in the 5' to 3' direction.

| Evolved | Strain | IC50 (μg/ml) | Std Err |
|---|---|---|---|
| 5 thy | 1 | 750 | 21 |
| | 2 | 720 | 24 |
| | 3 | 710 | 85 |
| | 4 | 910 | 11 |
| | 5 | 770 | 7.7 |
| | 6 | 790 | 7.3 |
| | 7 | 870 | 12 |
| | 8 | 840 | 10 |
| | 9 | 640 | 46 |
| | 10 | 830 | 81 |
| 10 thy | 1 | 890 | 63 |
| | 2 | 870 | 19 |
| | 3 | 1000 | 37 |
| | 4 | 970 | 25 |
| | 5 | 900 | 150 |
| | 6 | 1000 | 18 |
| | 7 | 830 | 120 |
| | 8 | 1100 | 37 |
| | 9 | 1000 | 140 |
| | 10 | 1000 | 31 |
| 50 thy | 1 | 1100 | 55 |
| | 2 | 780 | 29 |
| | 3 | 820 | 49 |
| | 4 | NA | NA |
| | 5 | NA | NA |
| | 6 | 1000 | 18 |
| | 7 | 820 | 77 |
| | 8 | 870 | 20 |
| | 9 | 760 | 65 |
| | 10 | NA | NA |

| Founder | Strain | IC50 (μg/ml) | Std Err |
|---|---|---|---|
| 5 thy | 1 | 0.91 | 0.036 |
| | 2 | 1.1 | 0.043 |
| 10 thy | 1 | 0.86 | 0.0071 |
| | 2 | 0.95 | 0.11 |
| 50 thy | 1 | 1.2 | 0.037 |
| | 2 | 1.3 | 0.091 |

| DHFR mut | Strain | IC50 (μg/ml) | Std Err |
|---|---|---|---|
| 5 thy | WT | 4.6 | 0.049 |
| | P21L | 41 | 0.5 |
| | W30R | 14 | 0.6 |
| | L28R | 370 | 4.9 |
| 50 thy | WT | 5.1 | 0.21 |
| | P21L | 40 | 0.72 |
| | W30R | 18 | 0.61 |
| | L28R | 420 | 8.9 |

| Recon | Strain | IC50 (μg/ml) | Std Err |
|---|---|---|---|
| 5 thy | WT/Δ25-26 | NA | NA |
| | P21L/Δ25-26 | 450 | 25 |
| | W30R/Δ25-26 | 580 | 25 |
| | L28RR/Δ25-26 | 1100 | 21 |
| 50 thy | WT/Δ25-26 | NA | NA |
| | P21L/Δ25-26 | 820 | 81 |
| | W30R/Δ25-26 | 1000 | 45 |
| | L28RR/Δ25-26 | >1800 | NA |

Table S6 **Trimethoprim resistance (IC50) for forward evolution strains.** In the leftmost panel, each strain is listed under the thymidine concentration used for both evolution and phenotyping. The top right panel contains IC50 measurements for unevolved founder strains in all four conditions. The remaining panels contain IC50 phenotypes of genetically engineered strains with either a single DHFR mutation, or pairs of DHFR/TYMS genotypes referred to as 'reconstitution' strains. These were measured in two thymidine concentrations, 5 and 50μg/ml. An estimate could not be obtained for most strains with a loss-of-function in TYMS paired with a wild-type DHFR (evolved strains 50thy-4,5,10; recon strain 1), since these grew slowly in all

TMP concentrations and without a sigmoidal dose-response. Standard error is calculated across triplicate measurements.

| Evolved | Strain | Coverage | Dispersion ($\sigma^2/\mu$) | Reads | Avg read length (BP) |
|---|---|---|---|---|---|
| 5 thy | 1 | 55 | 4.4 | 2.07E+06 | 126 |
| | 2 | 40 | 3.4 | 1.52E+06 | 125 |
| | 3 | 40 | 3.4 | 1.43E+06 | 131 |
| | 4 | 35 | 3.5 | 1.46E+06 | 114 |
| | 5 | 29 | 2.3 | 9.85E+05 | 138 |
| | 6 | 38 | 3.0 | 1.45E+06 | 122 |
| | 7 | 46 | 3.0 | 1.53E+06 | 142 |
| | 8 | 68 | 4.1 | 2.30E+06 | 137 |
| | 9 | 52 | 3.2 | 1.70E+06 | 140 |
| | 10 | 52 | 4.2 | 1.80E+06 | 134 |
| 10 thy | 1 | 32 | 3.4 | 1.13E+06 | 135 |
| | 2 | 34 | 3.3 | 1.16E+06 | 138 |
| | 3 | 34 | 3.4 | 1.19E+06 | 135 |
| | 4 | 48 | 3.9 | 1.67E+06 | 135 |
| | 5 | 42 | 3.6 | 1.46E+06 | 133 |
| | 6 | 31 | 2.9 | 1.12E+06 | 132 |
| | 7 | 29 | 2.9 | 1.03E+06 | 131 |
| | 8 | 25 | 2.6 | 8.43E+05 | 137 |
| | 9 | 31 | 2.9 | 1.07E+06 | 135 |
| | 10 | 41 | 3.4 | 1.44E+06 | 135 |
| 50 thy | 1 | 42 | 3.5 | 1.46E+06 | 133 |
| | 2 | 35 | 3.4 | 1.32E+06 | 126 |
| | 3 | 33 | 3.0 | 1.16E+06 | 134 |
| | 4 | 21 | 3.1 | 8.79E+05 | 116 |
| | 5 | 18 | 3.7 | 7.93E+05 | 115 |
| | 6 | 25 | 3.5 | 9.90E+05 | 121 |
| | 7 | 24 | 2.9 | 8.87E+05 | 125 |
| | 8 | 31 | 3.2 | 1.13E+06 | 130 |
| | 9 | 29 | 2.9 | 1.00E+06 | 136 |
| | 10 | 24 | 3.2 | 9.36E+05 | 126 |

| Founders | Strain | Coverage | Dispersion ($\sigma^2/\mu$) | Reads | Avg read length (BP) |
|---|---|---|---|---|---|
| 5 thy | 1 | 44 | 3.1 | 1.53E+06 | 134 |
| | 2 | 50 | 3.2 | 1.92E+06 | 122 |
| 10 thy | 1 | 36 | 2.2 | 1.32E+06 | 129 |
| | 2 | 40 | 2.5 | 1.45E+06 | 130 |
| 50 thy | 1 | 35 | 2.3 | 1.18E+06 | 137 |
| | 2 | 38 | 2.5 | 1.26E+06 | 138 |

Table S7 **Whole genome sequencing statistics**. Coverage is the average number of reads aligned to a particular position in the genome. Dispersion is the variance of read coverage normalized by the mean.

| Name | Mutation | | Strains | Annotation |
|---|---|---|---|---|
| folA | CCG to CTG | P21L | 10thy: 7 | dihydrofolate reductase |
| | CTC to CGC | L28R | 5thy: 1, 2, 3, 4, 5, 6, 7, 8, 9, 10; 50thy: 1, 6 | |
| | TGG to AGG | W30R | 10thy: 1, 2, 3, 4, 5, 6, 8, 9, 10; 50thy: 2, 3, 8, 9 | |
| thyA | IS1(+) +9bp | (627-635/795 nt) | 50thy: 2, 3, 8, 9 | thymidylate synthetase |
| | IS1(+) +9bp | (564-572/795 nt) | 50thy: 4, 5, 10 | |
| | Δ1 | (535/795 nt) | 10thy: 1, 2, 3, 4, 5, 6, 7, 8, 9, 10 | |
| | Δ6 | (525-530/795 nt) | 50thy: 7 | |
| | TGG to AGG | W133R | 50thy: 1, 6 | |
| | Δ6 | (64-69/795 nt) | 5thy: 1, 2, 3, 4, 5, 6, 7, 8, 9, 10 | |
| dusB | IS1(+) +10bp | (573-582/966 nt) | 10thy: 1, 2, 3, 4, 5, 6, 7, 8, 9, 10 | tRNA-dihydrouridine synthase B |
| | IS1(+) +8bp | (818-825/966 nt) | 5thy: 1, 2, 3, 4, 5, 6, 7, 8, 9, 10 | |
| cynR | AAT to AAA | N272K | 5thy: 10 | transcriptional activator of cyn operon; autorepressor |
| | AAA to GAA | K271E | 5thy: 6, 9, 10; 10thy: 4, 5, 6, 10 | |
| | TTG to TTT | L267F | 5thy: 3, 8, 10; 10thy: 1, 7, 8; 50thy: 6 | |
| ychE/oppA | Δ1199 | (+254/-485) | 5thy: 2, 4, 5; 10thy: 2, 3, 7, 8, 9, 10 | UPF0056 family inner membrane protein/oligopeptide ABC transporter periplasmic binding protein |
| gadX | GAT to GGT | D38G | 5thy: 5; 10thy: 2 | acid resistance regulon transcriptional activator; autoactivator |
| | GCG to TCG | A37S | 10thy: 2, 3, 5, 6, 7, 9, 10; 50thy: 8 | |
| otsB/araH | A to G | (-136/+31) | 10thy: 2, 3; 50thy: 3, 9 | trehalose-6-phosphate phosphatase, biosynthetic/L-arabinose ABC transporter permease |
| | C to A | (-142/+25) | 5thy: 3; 10thy: 2 | |
| | G to A | (-164/+3) | 10thy: 7; 50thy: 9 | |
| yfbL/yfbM | G to A | (+31/-72) | 5thy: 6, 7, 8; 10thy: 2, 5, 10 | putative M28A family peptidase/DUF1877 family protein |
| betI | TCC to CCC | S182P | 10thy: 1, 2, 4 | choline-inducible betIBA-betT divergent operon transcriptional repressor |
| | ACC to CCC | T178P | 10thy: 4 | |
| | GAT to GAA | D176E | 10thy: 1, 4, 5 | |
| | GAT to AAT | D176N | 10thy: 2, 4 | |
| ybaL | TAA to GAA | *559E | 10thy: 8; 50thy: 10 | inner membrane putative NAD(P)-binding transporter |
| | GTG to GGG | V555G | 5thy: 5, 6 | |
| cat/egfp | C to G | (+289/-204) | 5thy: 3, 8; 10thy: 4, 5 | chloramphenicol acetyltransferase/green fluorescent protein |

| | | | | |
|---|---|---|---|---|
| | C to G | (+299/-194) | 10thy: 5 | |
| fis | TCG to TAG | S30* | 50thy: 2, 3, 8, 9 | global DNA-binding transcriptional dual regulator |
| lacI | CCC to CCA | P332P | 10thy: 8 | lactose-inducible lac operon transcriptional repressor |
| | ACC to CCC | T329P | 5thy: 6; 10thy: 7, 8 | |
| chaA | ACC to CCC | T10P | 50thy: 7 | calcium/sodium:proton antiporter |
| | GTA to GAA | V8E | 10thy: 5 | |
| | CAA to AAA | Q5K | 10thy: 2 | |
| csrA | TAA to TAC | *62Y | 50thy: 4, 5, 10 | pleiotropic regulatory protein for carbon source metabolism |
| citG | ACC to CCC | T255P | 5thy: 2; 10thy: 3 | 2-(5''-triphosphoribosyl)-3'-dephosphocoenzyme-A synthase |
| ompF/asnS | G to T | (-529/+74) | 10thy: 2 | outer membrane porin 1a (Ia;b;F)/asparaginyl tRNA synthetase |
| | C to A | (-540/+63) | 10thy: 8 | |
| lpoB | CAA to CAC | Q38H | 5thy: 6; 50thy: 9 | OM lipoprotein stimulator of MrcB transpeptidase |
| sapA | ACC to CCC | T304P | 5thy: 2; 50thy: 4 | antimicrobial peptide transport ABC transporter periplasmic binding protein |
| yddE | CAA to CAC | Q14H | 50thy: 4, 5 | PhzC-PhzF family protein |
| | ACC to CCC | T12P | 50thy: 4 | |
| yghQ | GTG to GGG | V332G | 5thy: 7; 10thy: 10 | putative inner membrane polysaccharide flippase |
| | GGA to GGG | G323G | 10thy: 10 | |
| agaD | GGA to GGG | G120G | 10thy: 1 | N-acetylgalactosamine-specific enzyme IID component of PTS |
| | GCC to TCC | A126S | 10thy: 1, 3 | |
| rtcA | AGT to GGT | S215G | 10thy: 3; 50thy: 9 | RNA 3'-terminal phosphate cyclase |
| gntR | GAA to GGA | E147G | 10thy: 4, 5 | d-gluconate inducible gluconate regulon transcriptional repressor |
| | GTG to GGG | V146G | 10thy: 4 | |
| yiaK | ACC to CCC | T309P | 10thy: 7; 50thy: 6 | 2,3-diketo-L-gulonate reductase, NADH-dependent |
| | GAA to AAA | E313K | 10thy: 7 | |
| rrfB/murB | C to G | (+126/-175) | 10thy: 2, 10 | 5S ribosomal RNA of rrnB operon/UDP-N-acetylenolpyruvoylglucosamine reductase, FAD-binding |
| ampC | GTA to GGA | V48G | 10thy: 1, 8 | penicillin-binding protein; beta-lactamase, intrinsically weak |
| thrC | CTC to ATC | L3I | 50thy: 7 | L-threonine synthase |
| dapB/carA | T to A | (+301/-155) | 50thy: 6 | dihydrodipicolinate reductase/carbamoyl phosphate synthetase small subunit, glutamine amidotransferase |

| | | | | |
|---|---|---|---|---|
| paoC | CAA to AAA | Q72K | 50thy: 9 | PaoABC aldehyde oxidoreductase, Moco-containing subunit |
| acrR | IS1(+) +9bp | (320-328/648 nt) | 50thy: 7 | transcriptional repressor |
| ybdK | TGG to CGG | W263R | 5thy: 5 | weak gamma-glutamyl:cysteine ligase |
| dtpD/ybgI | T to A | (-84/-187) | 10thy: 3 | dipeptide and tripeptide permease D/NIF3 family metal-binding protein |
| ssuB | GGC to GGG | G44G | 10thy: 3, 8 | aliphatic sulfonate ABC transporter ATPase |
| | GTG to GGG | V43G | 50thy: 10 | |
| putP | GAT to GGT | D55G | 50thy: 3 | proline:sodium symporter |
| serX | A to G | (72/88 nt) | 10thy: 8 | tRNA-Ser |
| flgF | CAG to CGG | Q19R | 50thy: 10 | flagellar component of cell-proximal portion of basal-body rod |
| pabC | TAC to GAC | Y92D | 10thy: 3 | 4-amino-4-deoxychorismate lyase component of para-aminobenzoate synthase multienzyme complex |
| dadX | ACC to CCC | T284P | 5thy: 2 | alanine racemase, catabolic, PLP-binding |
| oppF | CCG to CAG | P273Q | 10thy: 3 | oligopeptide ABC transporter ATPase |
| uspF/ompN | G to A | (-108/+33) | 5thy: 7 | stress-induced protein, ATP-binding protein/outer membrane pore protein N, non-specific |
| yneM/dgcZ | G to A | (+75/+144) | 10thy: 3 | inner membrane-associated protein/diguanylate cyclase, zinc-sensing |
| yebV/yebW | G to T | (+26/-79) | 10thy: 7 | uncharacterized protein/uncharacterized protein |
| araH | CAA to AAA | Q322K | 10thy: 2 | L-arabinose ABC transporter permease |
| mntH | GTG to GGG | V313G | 10thy: 1 | manganese/divalent cation transporter |
| xapR | ATG to ATA | M176I | 5thy: 6 | transcriptional activator of xapAB |
| uraA | ATT to GTT | I311V | 50thy: 7 | uracil permease |
| relA | CAT to CAA | H518Q | 5thy: 3 | (p)ppGpp synthetase I/GTP pyrophosphokinase |
| ptrA | GAT to GAA | D38E | 10thy: 1 | protease III |
| | GAT to AAT | D38N | 10thy: 1 | |
| | CGT to CGA | R35R | 10thy: 1 | |
| rsmI | CAT to CAA | H235Q | 10thy: 6 | 16S rRNA C1402 2'-O-ribose methyltransferase, SAM-dependent |
| gltF/yhcA | Δ4 | (+90/-79) | 50thy: 7 | periplasmic protein/putative periplasmic chaperone protein |
| fis-yhdX | Δ9555 | | 50thy: 7 | fis, yhdJ, yhdU, acrS, acrE, acrF, yhdV, yhdW, yhdX |
| acrS | TAT to TTT | Y187F | 10thy: 3 | acrAB operon transcriptional repressor |
| secY | GTA to GGA | V274G | 10thy: 2 | preprotein translocase membrane subunit |
| xylF | GAA to AAA | E195K | 50thy: 6 | D-xylose transporter subunit |
| uhpT | GAA to GGA | E447G | 50thy: 8 | hexose phosphate transporter |

| | | | | |
|---|---|---|---|---|
| pstA | GGT to GGG | G112G | 50thy: 5 | phosphate ABC transporter permease |
| | ATT to GTT | I106V | 50thy: 5 | |
| pyrB | ACC to CCC | T54P | 50thy: 4 | aspartate carbamoyltransferase, catalytic subunit |

Table S8 **Annotated list of genes mutated during the forward evolution experiment**. Mutations identified in any of the founder strains are omitted. The first column indicates the affected gene; two names with a slash indicate neighboring genes to the affected intergenic region (ordered 5’ to 3’ along the sense strand). For proteins, both the codon change and amino acid change are included, synonymous mutations are omitted (an asterisk * indicates a stop codon). For intergenic mutations, the base change(s) and position relative to each neighboring gene are displayed. Insertion-sequence mediated changes are preceded with “IS#.” Annotations were pulled from the Breseq output.

| E. coli gene | Abbreviation | Name | Uniprot ID | Mutation | Description |
|---|---|---|---|---|---|
| *folA* | **DHFR** | dihydrofolate reductase | P0ABQ4 | coding | Catalyzes the reduction of THF to DHF, an essential reaction for de novo glycine and purine synthesis, and for DNA precursor synthesis. |
| *thyA* | **TYMS** | thymidylate synthase | P0A884 | coding | Catalyzes the reduction of dUMP to dTMP while utilizing 5,10-methylene THF as the methyl donor and reductant in the reaction, DHF as a by-product |
| *dusB* | **DUSB** | tRNA-dihydrouridine synthase B | P0ABT5 | coding | Catalyzes the synthesis of 5,6-dihydrouridine via the reduction of the C5-C6 double bond of uridine on target tRNA. |
| *cynR* | **CYNR** | HTH-type transcriptional regulator CynR | P27111 | coding | Positively regulates the cynTSX operon for cynate metabolism, and negatively regulates its own transcription. |
| *ychE* | **YHCE** | UPF0056 membrane protein YhcE | P25743 | intergenic (*/*oppA*) | Putative inner membrane protein. |
| *oppA* | **OPPA** | periplasmic oligopeptide-binding protein | P23843 | intergenic (*ychE*/*) | A component of the oligopeptide permease, a binding protein-dependent transport system. |
| *gadX* | **GADX** | HTH-type transcriptional regulator GadX | P37639 | coding | Positively regulates the expression of about fifteen genes involved in acid resistance such as gadA, gadB and gadC. Depending on the conditions (growth phase and medium), can repress gadW. |
| *otsB* | **TPP** | Trehalose-6-phosphate phosphatase | P31678 | intergenic (*/*araH*) | Removes the phosphate from trehalose 6-phosphate (Tre6P) to produce free trehalose. Also catalyzes the dephosphorylation of glucose-6-phosphate (Glu6P) and 2-deoxyglucose-6-phosphate (2dGlu6P). |
| *araH* | **ARAH** | L-arabinose transport system permease protein AraH | P0AE26 | intergenic (*otsB*/*) | Part of the binding-protein-dependent transport system for L-arabinose. Probably responsible for the translocation of the substrate across the membrane. |
| *yfbL* | **YFBL** | uncharacterized protein YfbL | P76482 | intergenic (*/*yfbM*) | |
| *yfbM* | **YFBM** | protein YfbM | P76483 | intergenic (*yfbL*/*) | |
| *betI* | **BETI** | HTH-type transcriptional regulator BetI | P17446 | coding | Repressor involved in the biosynthesis of the osmoprotectant glycine betaine. It represses transcription of the choline transporter BetT and the genes of BetAB involved in the synthesis of glycine betaine. |
| *ybaL* | **YBAL** | Putative cation/proton antiporter YbaL | P39830 | coding | Putative antiporter. |
| *cat* | **CAT** | chloramphenicol acetyltransferase | | intergenic (*/*eGFP*) | Chloramphenicol resistance marker introduced by phage transduciton. |
| *eGFP* | **EGFP** | enhanced green fluorescent protein | | intergenic (*cat*/*) | Enhanced green fluorescent protein introduced by phage transduction. |
| *fis* | **FIS** | DNA-binding protein Fis | P0A6R3 | coding | Activates ribosomal RNA transcription, as well other genes. Plays a direct role in upstream activation of rRNA promoters. Binds to hundreds of transcriptionally active and inactive AT-rich sites. |
| *lacI* | **LACI** | lactose operon repressor | P03023 | coding | Repressor of the lactose operon. Binds allolactose as an inducer. |
| *chaA* | **CHAA** | sodium-potassium/proton antiporter ChaA | P31801 | coding | Sodium exporter that functions mainly at alkaline pH. Can also function as a potassium/proton and calcium/proton antiporter at alkaline pH. |
| *csrA* | **CSR** | carbon storage regulator | P69913 | coding | A key translational regulator that binds mRNA to regulate translation initiation and/or mRNA stability, initially identified for its effects on central carbon metabolism. |
| *ttdR* | **TTDR** | HTH-type transcriptional activator TtdR | P45463 | intergenic *(*/ttdA*) | Positive regulator required for L-tartrate-dependent anaerobic growth on glycerol. Induces expression of the ttdA-ttdB-ygjE operon. |
| *ttdA* | **L-TTDα** | L(+)-tartrate dehydratase subunit alpha | P05847 | intergenic *(ttdR*/*) | Catalyzes the oxidation of (R,R)-tatrate to oxaloacetate. |

Table S9 **Functional annotations for commonly mutated genes.**

Each of the above genes was mutated at least 3 times across all conditions of the forward evolution experiment. Descriptions were paraphrased from UniProt.

Table S10 **Modular protein pairs identified by synteny**

*Sorted by distance from the diagonal. One representative gene name is given per COG. Pairs meeting the criteria in-pair > 1.0, out-pair < 0.5 are highlighted in orange.*

| COG 1 | gene 1 | COG 2 | gene 2 | physical interaction | shared metabolite | in-pair | out-pair |
|---|---|---|---|---|---|---|---|
| COG0103 | *rpsI* | COG0102 | *rplM* | in interaction | no shared intermediate | 3.81 | 0.82 |
| COG1271 | *cydA* | COG1294 | *cydB* | in interaction | no shared intermediate | 2.87 | 0.04 |
| COG1220 | *hslU* | COG5405 | *hslV* | in interaction | no shared intermediate | 3.14 | 0.55 |
| COG0719 | *sufD* | COG0396 | *sufC* | in interaction | no shared intermediate | 3.33 | 0.95 |
| COG0459 | *groL* | COG0234 | *groS* | in interaction | no shared intermediate | 2.56 | 0.23 |
| COG1108 | *znuB* | COG1121 | *znuC* | in interaction | no shared intermediate | 2.46 | 0.18 |
| COG1838 | *ttdB* | COG1951 | *fumA* | in interaction | shared int. | 2.47 | 0.23 |
| COG2884 | *ftsE* | COG2177 | *ftsX* | no interaction | no shared intermediate | 2.86 | 0.68 |
| COG0041 | *purE* | COG0026 | *purK* | no interaction | shared int. | 2.79 | 0.64 |
| COG3261 | *hyfG* | COG3260 | *hyfI* | in interaction | no shared intermediate | 2.52 | 0.40 |
| COG0074 | *yahF* | COG0045 | *sucC* | in interaction | shared int. | 2.55 | 0.43 |
| COG2025 | *fixB* | COG2086 | *fixA* | in interaction | no shared intermediate | 2.48 | 0.38 |
| COG0261 | *rplU* | COG0211 | *rpmA* | in interaction | no shared intermediate | 3.81 | 1.82 |
| COG0752 | *glyQ* | COG0751 | *glyS* | in interaction | shared int. | 2.67 | 0.82 |
| COG1918 | *feoA* | COG0370 | *feoB* | no interaction | no shared intermediate | 2.05 | 0.24 |
| COG1203 | *ygcB* | COG1518 | *ygbT* | no interaction | no shared intermediate | 2.00 | 0.23 |
| COG0048 | *rpsL* | COG0049 | *rpsG* | in interaction | no shared intermediate | 3.69 | 2.05 |
| COG0420 | *sbcD* | COG0419 | *sbcC* | in interaction | no shared intermediate | 1.80 | 0.17 |
| COG0208 | *nrdB* | COG0209 | *nrdA* | in interaction | shared int. | 1.79 | 0.18 |
| COG0184 | *rpsO* | COG1185 | *pnp* | no interaction | no shared intermediate | 2.89 | 1.43 |
| COG0052 | *rpsB* | COG0264 | *tsf* | no interaction | no shared intermediate | 3.51 | 2.12 |
| COG0725 | *modA* | COG4149 | *modB* | in interaction | no shared intermediate | 2.57 | 1.20 |
| COG0732 | *hsdS* | COG0286 | *hsdM* | no interaction | no shared intermediate | 1.67 | 0.32 |
| COG0262 | *folA* | COG0207 | *thyA* | no interaction | shared int. | 1.55 | 0.20 |
| COG1291 | *motA* | COG1360 | *motB* | in interaction | no shared intermediate | 2.31 | 1.01 |
| COG0505 | *carA* | COG0458 | *carB* | in interaction | shared int. | 1.68 | 0.39 |
| COG1702 | *ybeZ* | COG0319 | *ybeY* | no interaction | no shared intermediate | 2.80 | 1.51 |
| COG2104 | *thiS* | COG2022 | *thiG* | no interaction | no shared intermediate | 2.21 | 0.95 |
| COG1826 | *tatA* | COG0805 | *tatC* | in interaction | no shared intermediate | 1.36 | 0.10 |
| COG0004 | *amtB* | COG0347 | *glnK* | in interaction | no shared intermediate | 1.33 | 0.16 |
| COG1516 | *fliS* | COG1345 | *fliD* | no interaction | no shared intermediate | 2.71 | 1.54 |
| COG0066 | *leuD* | COG0065 | *leuC* | in interaction | shared int. | 2.28 | 1.16 |
| COG1843 | *flgD* | COG1749 | *flgE* | no interaction | no shared intermediate | 2.88 | 1.77 |
| COG1740 | *hyaA* | COG0374 | *hyaB* | in interaction | shared int. | 2.74 | 1.65 |
| COG0851 | *minE* | COG2894 | *minD* | in interaction | no shared intermediate | 3.77 | 2.68 |
| COG0168 | *trkG* | COG0569 | *kch* | no interaction | no shared intermediate | 1.17 | 0.08 |
| COG1585 | *ybbJ* | COG0330 | *qmcA* | no interaction | no shared intermediate | 1.12 | 0.07 |
| COG1077 | *mreB* | COG1792 | *mreC* | in interaction | no shared intermediate | 2.25 | 1.20 |
| COG0086 | *rpoC* | COG0085 | *rpoB* | in interaction | shared int. | 3.59 | 2.55 |
| COG0321 | *lipB* | COG0320 | *lipA* | no interaction | shared int. | 1.34 | 0.31 |
| COG0238 | *rpsR* | COG0360 | *rpsF* | in interaction | no shared intermediate | 3.41 | 2.37 |
| COG1925 | *ptsH* | COG1080 | *dhaM* | in interaction | no shared intermediate | 1.53 | 0.55 |
| COG0072 | *pheT* | COG0016 | *pheS* | in interaction | shared int. | 2.39 | 1.43 |
| COG1057 | *nadD* | COG0799 | *rsfS* | no interaction | no shared intermediate | 1.84 | 0.88 |

| COG 1 | gene 1 | COG 2 | gene 2 | physical interaction | shared metabolite | in-pair | out-pair |
|---|---|---|---|---|---|---|---|
| COG0280 | *eutD* | COG0282 | *ackA* | no interaction | no shared intermediate | 1.28 | 0.33 |
| COG0159 | *trpA* | COG0133 | *trpB* | in interaction | shared int. | 2.73 | 1.79 |
| COG1138 | *ccmF* | COG0755 | *ccmC* | in interaction | no shared intermediate | 1.17 | 0.23 |
| COG1921 | *selA* | COG3276 | *selB* | no interaction | no shared intermediate | 1.29 | 0.35 |
| COG0848 | *tolR* | COG0811 | *tolQ* | in interaction | no shared intermediate | 1.41 | 0.48 |
| COG0292 | *rplT* | COG0291 | *rpmI* | in interaction | no shared intermediate | 3.82 | 2.90 |
| COG0333 | *rpmF* | COG1399 | *yceD* | no interaction | no shared intermediate | 3.10 | 2.19 |
| COG1703 | *argK* | COG1884 | *scpA* | in interaction | no shared intermediate | 0.96 | 0.05 |
| COG0718 | *ybaB* | COG0353 | *recR* | no interaction | no shared intermediate | 2.60 | 1.71 |
| COG0742 | *rsmD* | COG0669 | *coaD* | no interaction | no shared intermediate | 1.45 | 0.57 |
| COG0194 | *gmk* | COG1561 | *yicC* | no interaction | no shared intermediate | 2.67 | 1.82 |
| COG0245 | *ispF* | COG1211 | *ispD* | no interaction | no shared intermediate | 1.36 | 0.52 |
| COG0263 | *proB* | COG0014 | *proA* | no interaction | shared int. | 1.88 | 1.03 |
| COG1825 | *rplY* | COG0193 | *pth* | no interaction | no shared intermediate | 2.29 | 1.47 |
| COG0689 | *rph* | COG0127 | *rdgB* | no interaction | no shared intermediate | 1.43 | 0.62 |
| COG2145 | *thiM* | COG0352 | *thiE* | no interaction | shared int. | 2.03 | 1.23 |
| COG0437 | *dmsB* | COG5557 | *hybB* | in interaction | no shared intermediate | 1.17 | 0.39 |
| COG0391 | *ybhK* | COG1660 | *yhbJ* | no interaction | no shared intermediate | 2.32 | 1.54 |
| COG0060 | *ileS* | COG0597 | *lspA* | no interaction | no shared intermediate | 1.17 | 0.40 |
| COG0248 | *ppx* | COG0855 | *ppk* | no interaction | shared int. | 1.09 | 0.33 |
| COG1923 | *hfq* | COG0324 | *miaA* | no interaction | no shared intermediate | 1.70 | 0.95 |
| COG0341 | *secF* | COG0342 | *secD* | in interaction | no shared intermediate | 3.27 | 2.52 |
| COG0439 | *accC* | COG0511 | *accB* | in interaction | no shared intermediate | 1.23 | 0.49 |
| COG0149 | *tpiA* | COG1314 | *secG* | no interaction | no shared intermediate | 1.57 | 0.84 |
| COG2087 | *cobU* | COG0368 | *cobS* | no interaction | shared int. | 2.19 | 1.47 |
| COG0528 | *pyrH* | COG0233 | *frr* | no interaction | no shared intermediate | 2.83 | 2.12 |
| COG0443 | *dnaK* | COG0576 | *grpE* | in interaction | no shared intermediate | 1.67 | 0.98 |
| COG0539 | *rpsA* | COG0283 | *cmk* | no interaction | no shared intermediate | 1.44 | 0.76 |
| COG0216 | *prfA* | COG2890 | *prmC* | no interaction | no shared intermediate | 1.56 | 0.88 |
| COG1137 | *lptB* | COG1934 | *lptA* | in interaction | no shared intermediate | 2.26 | 1.58 |
| COG2137 | *recX* | COG0468 | *recA* | in interaction | no shared intermediate | 1.41 | 0.74 |
| COG1975 | *paoD* | COG2068 | *mocA* | no interaction | no shared intermediate | 1.49 | 0.86 |
| COG0059 | *ilvC* | COG0440 | *ilvH* | no interaction | shared int. | 2.15 | 1.53 |
| COG2804 | *hofB* | COG1459 | *hofC* | in interaction | no shared intermediate | 1.69 | 1.08 |
| COG1177 | *potI* | COG1176 | *potH* | in interaction | no shared intermediate | 2.76 | 2.16 |
| COG0173 | *aspS* | COG0124 | *hisS* | no interaction | no shared intermediate | 1.00 | 0.39 |
| COG0407 | *hemE* | COG0276 | *hemH* | no interaction | no shared intermediate | 0.74 | 0.15 |
| COG1219 | *clpX* | COG0544 | *tig* | no interaction | no shared intermediate | 2.53 | 1.96 |
| COG1126 | *gltL* | COG0765 | *gltK* | in interaction | no shared intermediate | 1.67 | 1.11 |
| COG0636 | *atpE* | COG0356 | *atpB* | in interaction | no shared intermediate | 3.19 | 2.64 |
| COG1136 | *macB* | COG4591 | *lolC* | in interaction | no shared intermediate | 0.64 | 0.09 |
| COG0150 | *purM* | COG0299 | *purN* | in interaction | no shared intermediate | 1.44 | 0.89 |
| COG0297 | *glgA* | COG0448 | *glgC* | no interaction | shared int. | 1.14 | 0.59 |
| COG2009 | *sdhC* | COG0479 | *sdhB* | in interaction | no shared intermediate | 2.69 | 2.15 |

| COG 1 | gene 1 | COG 2 | gene 2 | physical interaction | shared metabolite | in-pair | out-pair |
|---|---|---|---|---|---|---|---|
| COG0409 | *hypD* | COG0298 | *hypC* | no interaction | no shared intermediate | 2.25 | 1.72 |
| COG0165 | *argH* | COG0137 | *argG* | no interaction | shared int. | 1.26 | 0.76 |
| COG1977 | *moaD* | COG0314 | *moaE* | in interaction | no shared intermediate | 0.95 | 0.46 |
| COG0470 | *holB* | COG0125 | *tmk* | no interaction | no shared intermediate | 0.96 | 0.47 |
| COG0594 | *rnpA* | COG0706 | *yidC* | no interaction | no shared intermediate | 2.91 | 2.43 |
| COG0044 | *allB* | COG0540 | *pyrB* | no interaction | no shared intermediate | 1.09 | 0.61 |
| COG0548 | *argA* | COG0002 | *argC* | no interaction | shared int. | 1.26 | 0.79 |
| COG0083 | *thrB* | COG0498 | *thrC* | no interaction | shared int. | 0.73 | 0.26 |
| COG0581 | *pstA* | COG0573 | *pstC* | in interaction | no shared intermediate | 3.18 | 2.71 |
| COG1674 | *ftsK* | COG2834 | *lolA* | no interaction | no shared intermediate | 0.76 | 0.30 |
| COG1043 | *lpxA* | COG0764 | *fabZ* | no interaction | shared int. | 1.71 | 1.26 |
| COG0363 | *nagB* | COG0364 | *zwf* | no interaction | no shared intermediate | 1.03 | 0.58 |
| COG0329 | *yagE* | COG0289 | *dapB* | no interaction | shared int. | 0.97 | 0.53 |
| COG0057 | *gapA* | COG0126 | *pgk* | no interaction | shared int. | 1.28 | 0.84 |
| COG2011 | *metI* | COG1135 | *metN* | in interaction | no shared intermediate | 2.92 | 2.49 |
| COG1385 | *rsmE* | COG2264 | *prmA* | no interaction | no shared intermediate | 0.92 | 0.49 |
| COG0802 | *tsaE* | COG1214 | *tsaB* | in interaction | no shared intermediate | 1.02 | 0.60 |
| COG0414 | *panC* | COG0853 | *panD* | no interaction | shared int. | 1.49 | 1.07 |
| COG1558 | *flgC* | COG1815 | *flgB* | no interaction | no shared intermediate | 3.63 | 3.21 |
| COG0055 | *atpD* | COG0355 | *atpC* | in interaction | no shared intermediate | 3.20 | 2.79 |
| COG0516 | *guaC* | COG0519 | *guaA* | no interaction | shared int. | 0.57 | 0.17 |
| COG0244 | *rplJ* | COG0222 | *rplL* | in interaction | no shared intermediate | 3.71 | 3.31 |
| COG1806 | *ppsR* | COG0574 | *ppsA* | no interaction | no shared intermediate | 1.16 | 0.77 |
| COG1127 | *mlaF* | COG0767 | *mlaE* | in interaction | no shared intermediate | 1.77 | 1.38 |
| COG0421 | *speE* | COG1586 | *speD* | no interaction | shared int. | 0.65 | 0.26 |
| COG1410 | *metH* | COG0685 | *metF* | no interaction | shared int. | 0.53 | 0.16 |
| COG0713 | *nuoK* | COG0839 | *nuoJ* | in interaction | shared int. | 3.16 | 2.79 |
| COG0155 | *cysI* | COG0175 | *cysD* | no interaction | shared int. | 0.89 | 0.53 |
| COG1932 | *serC* | COG0111 | *ghrA* | no interaction | shared int. | 0.58 | 0.23 |
| COG1116 | *ssuB* | COG0600 | *tauC* | in interaction | no shared intermediate | 1.85 | 1.51 |
| COG1120 | *fhuC* | COG0609 | *fhuB* | in interaction | no shared intermediate | 1.55 | 1.21 |
| COG1173 | *gsiD* | COG0601 | *gsiC* | in interaction | no shared intermediate | 1.26 | 0.93 |
| COG2127 | *clpS* | COG2360 | *aat* | no interaction | no shared intermediate | 0.83 | 0.50 |
| COG1438 | *argR* | COG0497 | *recN* | no interaction | no shared intermediate | 1.53 | 1.22 |
| COG0823 | *tolB* | COG1729 | *ybgF* | no interaction | no shared intermediate | 0.77 | 0.46 |
| COG1905 | *nuoE* | COG1894 | *nuoF* | in interaction | shared int. | 1.96 | 1.65 |
| COG0106 | *hisA* | COG0131 | *hisB* | no interaction | no shared intermediate | 2.07 | 1.77 |
| COG2001 | *mraZ* | COG0275 | *rsmH* | no interaction | no shared intermediate | 3.45 | 3.15 |
| COG0069 | *gltB* | COG0493 | *preT* | in interaction | shared int. | 0.53 | 0.23 |
| COG1195 | *recF* | COG0592 | *dnaN* | no interaction | no shared intermediate | 1.77 | 1.47 |
| COG0247 | *ykgE* | COG1139 | *ykgF* | no interaction | no shared intermediate | 0.45 | 0.16 |
| COG0375 | *hypA* | COG0378 | *hypB* | no interaction | no shared intermediate | 1.74 | 1.46 |
| COG0260 | *pepB* | COG0795 | *lptF* | no interaction | no shared intermediate | 0.37 | 0.10 |
| COG1319 | *paoB* | COG2080 | *paoA* | in interaction | no shared intermediate | 1.91 | 1.64 |
| COG0317 | *relA* | COG1490 | *dtd* | no interaction | no shared intermediate | 0.66 | 0.39 |
| COG1256 | *flgK* | COG1551 | *csrA* | no interaction | no shared intermediate | 1.96 | 1.70 |
| COG0304 | *fabF* | COG0236 | *acpP* | in interaction | no shared intermediate | 1.32 | 1.06 |
| COG0543 | *fre* | COG0167 | *pyrD* | no interaction | no shared intermediate | 0.74 | 0.49 |
| COG0550 | *topA* | COG0758 | *smf* | no interaction | no shared intermediate | 0.80 | 0.55 |
| COG0836 | *cpsB* | COG1089 | *gmd* | no interaction | shared int. | 0.59 | 0.35 |
| COG0701 | *yraQ* | COG0640 | *ygaV* | no interaction | no shared intermediate | 0.45 | 0.22 |
| COG1338 | *fliP* | COG1987 | *fliQ* | no interaction | no shared intermediate | 3.10 | 2.87 |
| COG0664 | *yaiV* | COG1151 | *hcp* | no interaction | no shared intermediate | 0.30 | 0.06 |
| COG0161 | *bioA* | COG0132 | *bioD* | no interaction | shared int. | 1.24 | 1.01 |
| COG2513 | *prpB* | COG0372 | *prpC* | no interaction | no shared intermediate | 0.33 | 0.12 |
| COG0801 | *folK* | COG1539 | *folX* | no interaction | shared int. | 0.99 | 0.78 |
| COG0228 | *rpsP* | COG0806 | *rimM* | no interaction | no shared intermediate | 3.05 | 2.85 |
| COG0128 | *aroA* | COG0287 | *tyrA* | no interaction | no shared intermediate | 0.97 | 0.76 |
| COG4775 | *bamA* | COG2825 | *skp* | no interaction | no shared intermediate | 1.47 | 1.26 |
| COG2255 | *ruvB* | COG0632 | *ruvA* | in interaction | no shared intermediate | 1.99 | 1.79 |
| COG0710 | *aroD* | COG0169 | *ydiB* | no interaction | shared int. | 0.81 | 0.61 |
| COG0838 | *nuoA* | COG0377 | *nuoB* | in interaction | shared int. | 2.70 | 2.50 |
| COG0643 | *cheA* | COG2201 | *cheB* | no interaction | no shared intermediate | 1.56 | 1.36 |
| COG1074 | *recB* | COG0507 | *recD* | in interaction | no shared intermediate | 0.30 | 0.11 |
| COG1937 | *frmR* | COG2217 | *copA* | no interaction | no shared intermediate | 0.29 | 0.11 |
| COG0691 | *smpB* | COG0557 | *rnr* | no interaction | no shared intermediate | 0.57 | 0.39 |
| COG0274 | *deoC* | COG0213 | *deoA* | no interaction | no shared intermediate | 0.59 | 0.41 |
| COG0112 | *glyA* | COG0698 | *rpiB* | no interaction | no shared intermediate | 0.51 | 0.34 |
| COG0237 | *coaE* | COG0749 | *polA* | no interaction | no shared intermediate | 0.55 | 0.38 |
| COG0568 | *rpoS* | COG0358 | *dnaG* | no interaction | no shared intermediate | 0.92 | 0.75 |
| COG0195 | *nusA* | COG0779 | *rimP* | no interaction | no shared intermediate | 3.51 | 3.35 |
| COG2919 | *ftsB* | COG0148 | *eno* | no interaction | no shared intermediate | 0.86 | 0.70 |
| COG0736 | *acpS* | COG0063 | *nnr* | no interaction | no shared intermediate | 0.74 | 0.60 |
| COG0279 | *gmhA* | COG0241 | *gmhB* | no interaction | no shared intermediate | 0.83 | 0.68 |
| COG0080 | *rplK* | COG0081 | *rplA* | in interaction | no shared intermediate | 3.52 | 3.38 |
| COG0524 | *gsk* | COG0800 | *eda* | no interaction | shared int. | 0.33 | 0.20 |
| COG0825 | *accA* | COG0777 | *accD* | in interaction | shared int. | 0.76 | 0.63 |
| COG1587 | *hemD* | COG0181 | *hemC* | no interaction | shared int. | 1.41 | 1.29 |
| COG3383 | *fdhF* | COG1526 | *fdhD* | no interaction | no shared intermediate | 0.51 | 0.39 |
| COG0084 | *ycfH* | COG0143 | *metG* | no interaction | no shared intermediate | 0.50 | 0.38 |
| COG0554 | *glpK* | COG0578 | *glpA* | no interaction | shared int. | 1.41 | 1.29 |
| COG2878 | *rsxB* | COG0177 | *nth* | no interaction | no shared intermediate | 0.23 | 0.12 |
| COG0445 | *mnmG* | COG0357 | *rsmG* | no interaction | no shared intermediate | 1.22 | 1.11 |
| COG2877 | *kdsA* | COG0504 | *pyrG* | no interaction | no shared intermediate | 0.83 | 0.72 |
| COG1596 | *gfcE* | COG2148 | *wcaJ* | no interaction | no shared intermediate | 0.52 | 0.41 |
| COG0583 | *nhaR* | COG2855 | *yeiH* | no interaction | no shared intermediate | 0.14 | 0.03 |
| COG0508 | *aceF* | COG0567 | *sucA* | in interaction | shared int. | 0.58 | 0.47 |
| COG0325 | *yggS* | COG1496 | *yfiH* | no interaction | no shared intermediate | 0.50 | 0.40 |
| COG0337 | *aroB* | COG0703 | *aroL* | no interaction | no shared intermediate | 1.01 | 0.91 |
| COG1995 | *pdxA* | COG0030 | *rsmA* | no interaction | no shared intermediate | 0.54 | 0.44 |
| COG0043 | *ubiD* | COG0163 | *ubiX* | in interaction | no shared intermediate | 0.28 | 0.19 |
| COG0381 | *wecB* | COG0677 | *wecC* | no interaction | shared int. | 0.50 | 0.41 |

| COG 1 | gene 1 | COG 2 | gene 2 | physical interaction | shared metabolite | in-pair | out-pair |
|---|---|---|---|---|---|---|---|
| COG0272 | *ligA* | COG0210 | *helD* | no interaction | no shared intermediate | 0.19 | 0.09 |
| COG0054 | *ribE* | COG0307 | *ribC* | no interaction | shared int. | 1.70 | 1.61 |
| COG0242 | *def* | COG0223 | *arnA* | no interaction | no shared intermediate | 1.04 | 0.94 |
| COG1129 | *lsrA* | COG1172 | *lsrC* | in interaction | no shared intermediate | 1.51 | 1.42 |
| COG1488 | *pncB* | COG1335 | *ycaC* | no interaction | shared int. | 0.21 | 0.12 |
| COG0746 | *mobA* | COG1763 | *mobB* | no interaction | no shared intermediate | 0.42 | 0.34 |
| COG0328 | *rnhA* | COG2334 | *rdoA* | no interaction | no shared intermediate | 0.18 | 0.10 |
| COG1003 | *gcvP* | COG0509 | *gcvH* | in interaction | no shared intermediate | 1.52 | 1.43 |
| COG0845 | *acrA* | COG0841 | *acrB* | in interaction | no shared intermediate | 0.35 | 0.27 |
| COG0227 | *rpmB* | COG1200 | *recG* | no interaction | no shared intermediate | 0.65 | 0.57 |
| COG1886 | *fliN* | COG1868 | *fliM* | in interaction | no shared intermediate | 2.26 | 2.18 |
| COG0395 | *ycjP* | COG1175 | *ycjO* | in interaction | no shared intermediate | 1.03 | 0.95 |
| COG1045 | *wcaB* | COG0031 | *cysK* | in interaction | shared int. | 0.32 | 0.24 |
| COG0178 | *uvrA* | COG0556 | *uvrB* | in interaction | no shared intermediate | 0.49 | 0.42 |
| COG0203 | *rplQ* | COG0100 | *rpsK* | in interaction | no shared intermediate | 3.85 | 3.77 |
| COG0046 | *purL* | COG0152 | *purC* | no interaction | no shared intermediate | 0.82 | 0.75 |
| COG0164 | *rnhB* | COG0792 | *yraN* | no interaction | no shared intermediate | 1.18 | 1.11 |
| COG3956 | *mazG* | COG1188 | *hslR* | no interaction | no shared intermediate | 0.78 | 0.71 |
| COG1162 | *rsgA* | COG0036 | *rpe* | no interaction | no shared intermediate | 0.69 | 0.63 |
| COG1250 | *paaH* | COG0183 | *paaJ* | in interaction | shared int. | 0.30 | 0.24 |
| COG0782 | *rnk* | COG1190 | *lysS* | no interaction | no shared intermediate | 0.42 | 0.36 |
| COG0602 | *queE* | COG0603 | *queC* | no interaction | shared int. | 0.98 | 0.92 |
| COG1841 | *rpmD* | COG0200 | *rplO* | in interaction | no shared intermediate | 3.73 | 3.67 |
| COG0712 | *atpH* | COG0056 | *atpA* | in interaction | no shared intermediate | 3.27 | 3.22 |
| COG0527 | *thrA* | COG0136 | *usg* | no interaction | shared int. | 0.43 | 0.38 |
| COG1570 | *xseA* | COG1722 | *xseB* | in interaction | no shared intermediate | 1.85 | 1.80 |
| COG0465 | *ftsH* | COG0037 | *tilS* | no interaction | no shared intermediate | 0.77 | 0.72 |
| COG0088 | *rplD* | COG0089 | *rplW* | in interaction | no shared intermediate | 3.86 | 3.81 |
| COG1622 | *cyoA* | COG0843 | *cyoB* | in interaction | shared int. | 2.04 | 1.99 |
| COG0229 | *msrB* | COG0225 | *msrA* | no interaction | shared int. | 0.11 | 0.06 |
| COG0040 | *hisG* | COG0141 | *hisD* | no interaction | no shared intermediate | 1.15 | 1.10 |
| COG0092 | *rpsC* | COG0255 | *rpmC* | in interaction | no shared intermediate | 3.86 | 3.82 |
| COG1597 | *yegS* | COG1409 | *cpdA* | no interaction | no shared intermediate | 0.07 | 0.03 |
| COG1070 | *lsrK* | COG0235 | *araD* | no interaction | shared int. | 0.15 | 0.11 |
| COG0547 | *ybiB* | COG0134 | *trpC* | no interaction | shared int. | 1.83 | 1.79 |
| COG0256 | *rplR* | COG0096 | *rpsH* | in interaction | no shared intermediate | 3.72 | 3.67 |
| COG1131 | *yadG* | COG0842 | *ybhR* | no interaction | no shared intermediate | 0.32 | 0.28 |
| COG4177 | *livM* | COG0559 | *livH* | in interaction | no shared intermediate | 1.70 | 1.66 |
| COG0787 | *dadX* | COG2337 | *mazF* | no interaction | no shared intermediate | 0.40 | 0.37 |

| COG 1 | gene 1 | COG 2 | gene 2 | physical interaction | shared metabolite | in-pair | out-pair |
|---|---|---|---|---|---|---|---|
| COG0481 | *lepA* | COG0635 | *yggW* | no interaction | no shared intermediate | 0.49 | 0.45 |
| COG1555 | *ybaV* | COG2333 | *ycaI* | no interaction | no shared intermediate | 0.79 | 0.76 |
| COG1317 | *fliH* | COG1766 | *fliF* | no interaction | no shared intermediate | 3.18 | 3.15 |
| COG0743 | *dxr* | COG0575 | *cdsA* | no interaction | no shared intermediate | 2.37 | 2.34 |
| COG0187 | *parE* | COG0188 | *gyrA* | in interaction | no shared intermediate | 1.01 | 0.98 |
| COG0659 | *dauA* | COG0288 | *can* | no interaction | no shared intermediate | 0.06 | 0.03 |
| COG0461 | *pyrE* | COG0284 | *pyrF* | in interaction | shared int. | 0.63 | 0.61 |
| COG0469 | *pykF* | COG0205 | *pfkA* | no interaction | shared int. | 0.31 | 0.29 |
| COG1559 | *yceG* | COG0816 | *yqgF* | no interaction | no shared intermediate | 0.92 | 0.90 |
| COG1576 | *rlmH* | COG1235 | *phnP* | no interaction | no shared intermediate | 0.40 | 0.38 |
| COG1575 | *menA* | COG0318 | *caiC* | no interaction | no shared intermediate | 0.11 | 0.09 |
| COG1252 | *ndh* | COG1477 | *apbE* | no interaction | no shared intermediate | 0.09 | 0.07 |
| COG1528 | *ftnB* | COG0450 | *ahpC* | no interaction | no shared intermediate | 0.09 | 0.08 |
| COG1251 | *norW* | COG2223 | *lacY* | no interaction | no shared intermediate | 0.15 | 0.13 |
| COG1088 | *rfbB* | COG1898 | *rfbC* | no interaction | shared int. | 2.00 | 1.98 |
| COG1328 | *nrdD* | COG1180 | *ybiY* | no interaction | no shared intermediate | 0.41 | 0.39 |
| COG0709 | *selD* | COG0425 | *yedF* | no interaction | no shared intermediate | 0.37 | 0.35 |
| COG1519 | *waaA* | COG1663 | *lpxK* | no interaction | shared int. | 0.45 | 0.44 |
| COG0157 | *nadC* | COG0379 | *nadA* | no interaction | shared int. | 1.30 | 1.29 |
| COG0863 | *yhdJ* | COG1194 | *mutY* | no interaction | no shared intermediate | 0.12 | 0.10 |
| COG1092 | *rlmI* | COG2606 | *ybaK* | no interaction | no shared intermediate | 0.05 | 0.03 |
| COG1902 | *nemA* | COG2141 | *ssuD* | no interaction | no shared intermediate | 0.04 | 0.02 |
| COG0017 | *asnS* | COG0116 | *rlmL* | no interaction | no shared intermediate | 0.10 | 0.09 |
| COG0090 | *rplB* | COG0185 | *rpsS* | in interaction | no shared intermediate | 3.58 | 3.57 |
| COG0655 | *wrbA* | COG1733 | *ytfH* | no interaction | no shared intermediate | 0.04 | 0.03 |
| COG0122 | *alkA* | COG0350 | *ogt* | no interaction | no shared intermediate | 0.10 | 0.09 |
| COG1752 | *rssA* | COG3264 | *mscK* | no interaction | no shared intermediate | 0.04 | 0.03 |
| COG1607 | *yciA* | COG0604 | *acuI* | no interaction | no shared intermediate | 0.04 | 0.03 |
| COG1381 | *recO* | COG1159 | *era* | no interaction | no shared intermediate | 1.21 | 1.20 |
| COG1171 | *ygeX* | COG0120 | *rpiA* | no interaction | no shared intermediate | 0.09 | 0.09 |
| COG0038 | *clcA* | COG0025 | *yjcE* | no interaction | no shared intermediate | 0.03 | 0.02 |
| COG4108 | *prfC* | COG0791 | *yaeF* | no interaction | no shared intermediate | 0.05 | 0.04 |
| COG0657 | *aes* | COG4221 | *ydfG* | no interaction | no shared intermediate | 0.03 | 0.02 |
| COG1004 | *ugd* | COG1210 | *galU* | no interaction | shared int. | 0.31 | 0.30 |
| COG1015 | *yhfW* | COG0180 | *trpS* | no interaction | no shared intermediate | 0.16 | 0.16 |
| COG1055 | *arsB* | COG0598 | *ydaN* | no interaction | no shared intermediate | 0.03 | 0.03 |
| COG2110 | *ymdB* | COG0705 | *glpG* | no interaction | no shared intermediate | 0.04 | 0.04 |
| COG2896 | *moaA* | COG0315 | *moaC* | in interaction | shared int. | 0.66 | 0.66 |
| COG1012 | *betB* | COG0160 | *puuE* | no interaction | shared int. | 0.03 | 0.03 |
